# Chromatin Landscape of Cancer Cell Lines Identifies Enhancer Subtypes

**DOI:** 10.64898/2026.08.01.741526

**Authors:** Mahinur Mattohti, Emre Arslan, Ayush T. Raman, Christopher Terranova, Zhiyi Liu, Elias Orouji, Sanjana Srinivasan, Veena Kochat, Ming Tang, Samirkumar B. Amin, Jonathan Schulz, Neha S. Samant, Anand K. Singh, Emmanuel Martinez-Ledesma, Sharmistha Sarkar, Marcus Coyle, Christopher Bristow, Faye M. Johnson, Curtis R. Pickering, Keila E. Torres, Jeffery N. Myers, Kunal Rai

## Abstract

Epigenetic aberrations are a hallmark of cancer; however, systematic chromatin state maps of cancer cells are unavailable. We generated and analyzed 803 histone mark profiles in 142 cancer cell lines and 114 human tumors belonging to 9 solid tumor types. Irrespective of their cell-of-origin, cancer cells segregate from normal tissues based on their enhancer patterns, suggesting enhancer deregulation is a fundamental epigenetic feature in cancer. Enhancer based clustering defined 5 distinct subgroups of cancer cells (EpiC1-5) with unique developmental trajectories, molecular features and dependencies. Importantly, we define a set of core TFs that are critical for EpiC-specific enhancer patterns and survival. Notably, EpiC4 represented a predominantly epigenetic, pan-cancer subtype that displays poor survival, activation and dependence on a FN1-CAV1-SRC-PI3K-AKT signaling network. Together, these data uncover enhancer heterogeneity in pan-cancer systems with identification of a novel enhancer-based subtype and identify potential new therapeutic targets associated with unique epigenetic features.

## INTRODUCTION

Epigenetic elements, such as DNA methylation and histone modifications, regulate the transcriptional output of neighboring genes^1,2^. These elements and their regulators play important roles in ensuring the normal development of various organisms. In the past few years, we have begun to understand the genomic landscape of histone modifications in healthy human tissues through massive efforts such as the Roadmap^3^ and ENCODE^4^ consortiums which have comprehensively profiled chromatin landscape, open chromatin regions and DNA methylation in normal healthy tissues and a variety of cell lines. Cancer genomic studies have identified somatic mutations, copy number alterations, and translocations in epigenetic regulators across many cancers^5^. These include members of the DNMT/TET, MLL, and SWI/SNF families, which regulate DNA methylation, histone modifications, and chromatin remodeling activities, respectively^6,7^. Therefore, it is important to understand the genome-wide landscape of these chromatin modifications and accessible regions in cancers. TCGA studies have defined the DNA methylation landscape across many human tumors and a recent study profiled accessible chromatin regions with use of ATAC-Seq in several hundred human tumors^8^. Aberrant activation of enhancers has been identified as a key feature of ependymoma, medulloblastoma and pediatric gliomas tumor types^9,10^. Importantly, however, other epigenetic elements have been shown to play roles in cancer progression as well. Examples include reprogramming of H3K9me3-marked heterochromatin regions in pancreatic cancer^11^, impaired H3K36 methylation in head and neck cancers^12^ and defects in H3K27me2/me3-marked polycomb elements in glioma^13^.

Established cancer cell lines are valuable *in vitro* model systems that are widely used in cancer research and drug discovery. However, we do not have sufficient understanding of aberrations in the chromatin states of these lines which can be a useful resource for identifying fundamental traits of the cancers as well as for correlations with functional studies in these model systems. Histone modification maps are used to define the gene regulatory landscape^14^. Basic unit of chromatin is a nucleosome that gets heavily modified. We define “chromatin state” as specific combinatorial pattern of the detectable chromatin modifications and consider this a basic epigenetic feature of a genomic locus. It is the specific patterns of modifications on a certain nucleosome that dictate its impact on the transcriptional output of the associated locus and its vicinity^15^. For example, the presence of H3K27me3 is associated with a transcriptionally repressive environment^16^, whereas H3K4me3 is associated with transcriptionally active promoters^17,18^. Additionally, H3K4me1 and H3K27Ac dually modified nucleosomes are present at enhancer elements^19^, whereas the presence of H3K79me2 or H3K36me3 coincides with transcribed regions^20^. In this manuscript, we define genome-wide profiles of these histone modification marks and their combinatorial patterns in 142 cell lines used by Cancer Cell Line Encyclopedia (CCLE)^21^ and Sanger Cell Line Project^22^. Combined with the richness of diverse, orthogonal data types in the CCLE and other projects, the chromatin state landscape of these cancer cell lines provides a unique and important layer to understand the regulation of cancer gene transcription that ultimately affect cancer prognosis and therapy.

## RESULTS

### Chromatin State Dynamics Across 142 Cancer Cell Lines

We generated chromatin state maps in 142 established cancer lines (from the CCLE, Sanger and MD Anderson Cancer Center^21,23,24^ encompassing 9 tumor types (**Fig. 1A, Extended Data Fig. 1A** and **Table S1)** by profiling 6 reference histone marks previously used by NIH Roadmap Epigenomics project^3^ and ENCODE^4^: H3K4me3 for Pol II-bound promoters^15,16^, H3K4me1 for enhancers^19^, H3K27ac for active regions^19^, H3K79me2 for transcribed regions^20^, H3K27me3 for polycomb repressed regions^16^, and H3K9me3 for heterochromatin. At the same time, we generated or analyzed published H3K27ac profiles from 114 tumors from 6 different tumor types [BRCA (n= 10)^25^, COAD (n = 34)^26^, GBM (n = 22), OVCA (n = 20)^27^, SKCM (n = 20)^28^, PRAD (n =8)]. Several sets of additional genomics datasets are available for the CCLE/Sanger cell lines and a subset of human tumors (profiled by the TCGA), such as somatic mutations, mRNA expression, copy number alterations, drug response, CRISPR/Cas9 and RNAi based functional genomic data (projects Achilles/DRIVE), DNA methylation and proteomic (Reverse Phase Protein Array) datasets^29^ (**Fig. 1B**).

**Fig. 1:**
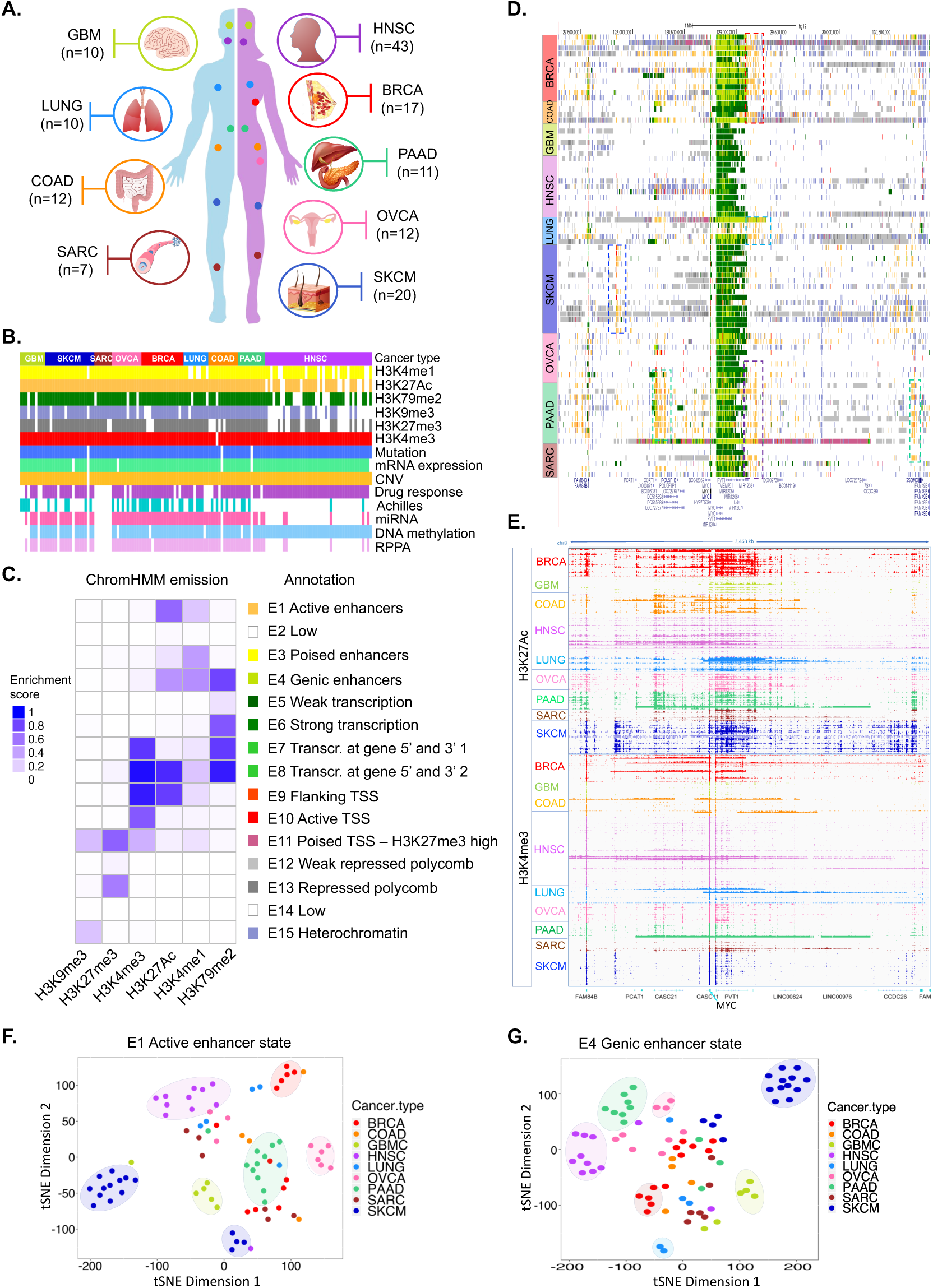
Comprehensive epigenome profiling of pan-cancer cell lines identifies diverse chromatin landscapes. **A.** Schematic illustrating the tissue of origin and number of cancer cell lines used for histone modification profiling in this study. See Table S1 for a full list of cell line names and other annotations. **B.** Schematic illustrating the six different histone modifications ChIP-seq data profiled in this study and other available data sets for each of the cancer cell lines. **C.** Emission probabilities of the 15-state ChromHMM model based on ChIP-seq profiles of six histone marks (shown on X-axis). Each row represents one chromatin state and each column corresponds to one chromatin mark. The intensity of the color in each cell reflects the frequency of occurrence of that mark in the corresponding chromatin state on a scale of 0 (white) to 1 (blue). States were manually grouped and given candidate annotations on the right. **D.** Chromatin state annotations across 80 cancer cell lines in an ∼3.5-Mb region near to the MYC gene on chromosome 8. Promoters are primarily constitutive (green vertical lines), whereas enhancers are highly dynamic (dispersed yellow regions). **E.** Genome browser view of ChIP-Seq signal tracks for H3K7AC(n=125) and H3K4me3(n=142) surrounding MYC gene region shown in panel 1D. **F.** Unsupervised t-SNE on the top 50 principal components for the 10,000 most variable active enhancer state regions across all cancer types. Each dot represents a given sample. Colors represents the cancer type shown above the plot. **G.** Unsupervised t-SNE on the top 50 principal components for the 10,000 most variable poised TSS – H3K27me3 high state regions across all cancer types. Each dot represents a given sample. Colors represents the cancer type shown above the plot.

To determine combinatorial patterns of the chromatin modifications that associate with biological functions and transcriptional outputs of associated genes^15^, we defined chromatin state maps using a 15-state model derived from ChromHMM algorithm^30^ (**Fig. 1C**) which consisted of the repertoire of chromatin states including promoters(E9, E10, E11), transcribed states (E5, E6, E7, E8), enhancers (E1, E3, E4), polycomb (E12, E13) and heterochromatin repression (E15) as well as poised states (E11) (**Fig. 1C**). Annotation of the chromatin states in a single track per cell line or of histone modifications is useful for the visualization of regulatory elements around cancer regulators. Notably, we observed marked enhancer gains unique to particular cancer subtypes around the MYC locus (**Fig. 1D**-**1E**) indicating different enhancers may be responsible for regulating the same oncogene^31^. Dimensionality reduction using tSNE technique suggested that the inherent active enhancer state best segregates cancer cells according to their histological subtypes (**Fig. 1F-1G** and **Extended Data Fig. 1B-1H**)^3^.

Importantly, cancer cell lines (n = 125) and normal samples (n = 93, from Roadmap/ENCODE) showed distinct clustering based on H3K27ac patterns irrespective of their tissue-of-origin (**Fig. 2A** and **Extended Data Fig. 2A**) suggesting enhancer states of cancers are inherently distinct from their corresponding normal. Since cancer cell lines are cultured in vitro for several years, we asked if the enhancer profiles from cell lines were present in human tumor samples. H3K27ac peaks from cancer cell lines of 5 different cancer types with showed significant (53-77%) overlaps with those derived from corresponding tumors (n = 105) [BRCA (n= 10)^25^, COAD (n = 34)^26^, GBM (n = 22), OVCA (n = 20)^27^, SKCM (n = 20)^28^] **(Fig. 2B),** similarly it showed ∼60% overlap with open chromatin regions from TCGA ATAC-Seq data (562,709 peaks)^8^ (**Extended Data Fig. 2B**). We identified 67,422 cancer-specific H3K27ac-marked enhancers (**Fig. 2C** and **Table S2**), only 21% of 67,422 peaks were shared among all different cancer types and the rest were distributed uniquely between cancer types (**Fig. 2D** and **Table S2),** with the highest representation in BRCA. The most intense enhancer peaks were observed in LUNG and the lowest intensity peaks were observed in HNSC and PAAD (**Fig. 2E** and **Extended Data Fig. 2C**). Examples of unique enhancer peaks included AXIN2 in colon cancer (**Fig. 2F, Extended Data Fig. 2D-2E** and **Table S2)** whereas CCND1 and ◻-catenin were present in a pan-cancer manner (**Extended Data Fig. 2F-2G**). The H3K27ac peaks were also validated in corresponding tumor samples (**Figures 2G** and **Extended Data Fig 2H-2I**). Hi-ChIP data showed specific interactions between these distal enhancers with their regulated gene promoters and robust association with increased transcription of the gene (**Fig. 2F** and **Extended Data Fig. 2J-2M**). Overlaps of cancer-specific enhancers with those in early developmental stages (ESCs, germ layers or their tissue-of-origin) suggested that while a substantial fraction likely arose due to re-activation of developmental enhancers, a higher number were gained in a *de novo* manner (**Fig. 2H**). De novo enhancers marked hyperactivated hallmark cancer signaling events whereas re-activated enhancers regulated pathways involved in cellular de-differentiation and patterning (**Fig. 2I, Extended Data Fig. 3A-3B** and **Table S3**). Integration of published GWAS data with cancer-specific enhancer calls identified enhancer SNPs that affect multiple enhancers of *bona fide* tumor suppressors and oncogenes including those around TERT gene promoter in breast cancers, lung and pancreatic adenocarcinomas as well as those around SIRT5^32^ in breast cancers (**Extended Data Fig. 3C-3D** and **Table S4**).

**Fig. 2:**
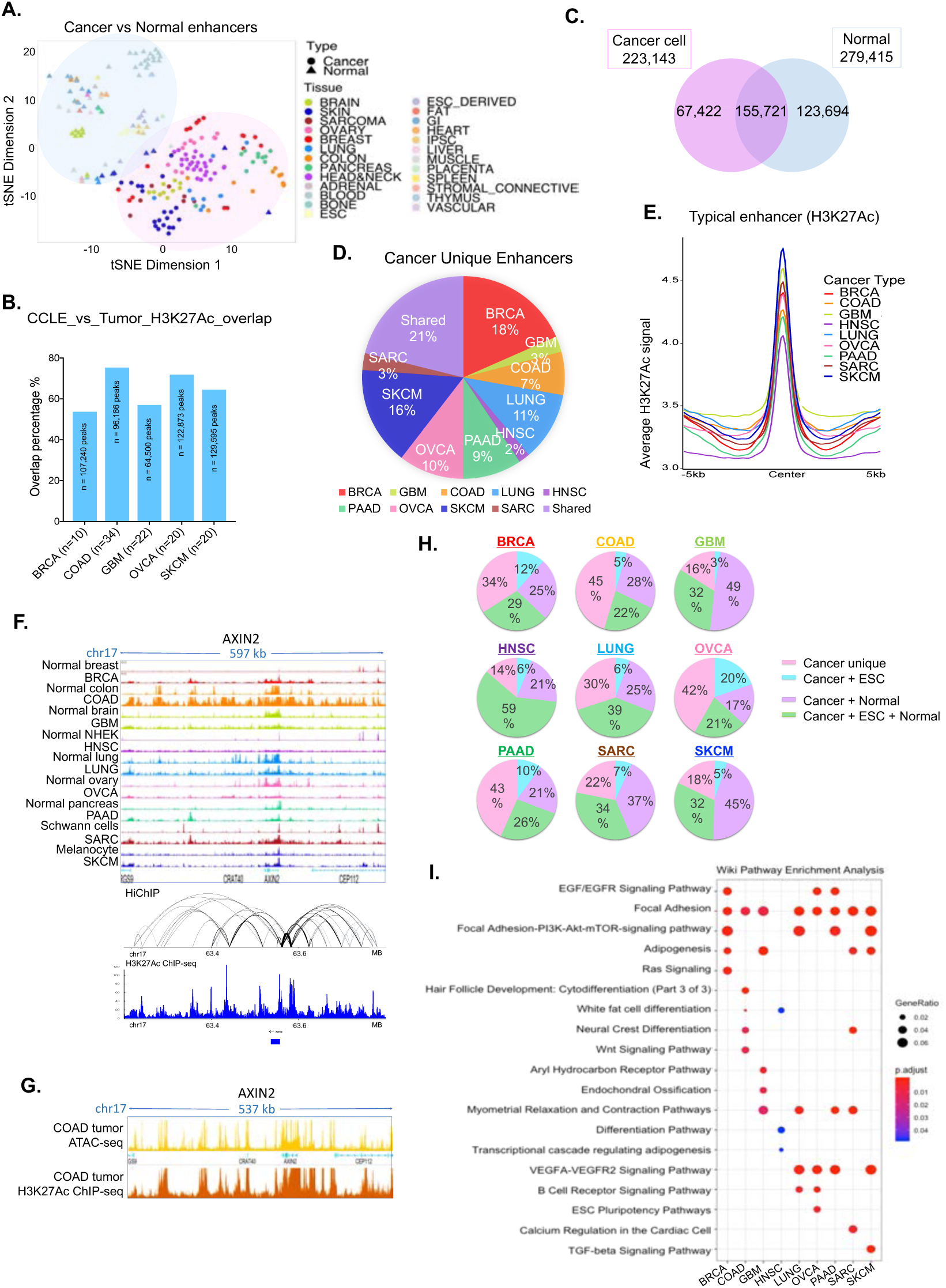
Cancer cells require cancer- and tissue-specific enhancer reprogramming during cancer development. **A.** Unsupervised t-SNE on the top 50 principal components for the 10,000 most variable enhancer peaks for 125 cancer cell lines and 93 Roadmap normal tissues. Circles represents cancer cell lines; triangles represent normal samples. Colors represents the tissue of origin shown above the plot. **B.** Bar plot showing the percentage of overlaps between H3K27Ac enhancer peaks from cancer cell lines and matched tumors. **C.** Venn diagram showing the overlap of all enhancer peaks outside +/-2.5kb TSS regions between cancer cell lines and normal samples to identify the cancer-unique or shared enhancers. **D.** Pie chart showing the percent overlap of all cancer-unique enhancer peaks to those present in each histological cancer types. **E.** Average intensity curves of ChIP-Seq reads (RPKM) for H3K27ac at typical enhancer regions. Enhancers are shown in a 10kb window centered on the middle of the enhancer in each cancer type. **F.** Genome browser view of H3K27Ac ChIP-Seq signal track(top) surrounding the AXIN2 gene as an example of colon cancer unique enhancers. Bottom panels show enhancer-promoter interaction loops for AXIN2 as derived from the H3K27Ac Hi-ChIP data in SW620 cells. **G.** Genome browser view of colon cancer TCGA ATAC-seq signal track(top) and H3K27Ac ChIP-Seq signal track(bottom) surrounding the AXIN2 gene. **H.** Pie chart showing the overlap ratios of H3K27ac peaks in each histological cancer type with normal or ESCs peaks to identify cancer-unique enhancer gains (*de novo*) (pink), memory enhancer re-gain from ESC (blue), cancer and normal common enhancers (purple), and consistent enhancers through development (green). **I.** Dot plot showing significantly enriched pathways for each cancer type *de novo* gain enhancers during development. Dot size represents gene ratio, and colors represents adjusted p-values.

### Enhancers Define Specific Clusters for Cancer Lines

Consensus Non-Negative Matrix Factorization (NMF) clustering on enhancer data from 124 cancer cell lines identified 5 distinct clusters of cancer cell lines, named as EpiC1 through EpiC5 (**Fig. 3A-3B**, **Extended Data Fig. 4A** and **Table S5**). The first three of these clusters correlated tightly based on their developmental origin including EpiC1 for melanoma, EpiC2 for breast and ovarian cancers and EpiC3 for colorectal and pancreatic cancers (**Fig. 3C)**. Cluster 4 (EpiC4) and Cluster 5 (EpiC5) were distinct in that they harbored cell lines from multiple different tissue types and were likely similar due to their molecular phenotypes (**Fig. 3C)**. NMF clustering based on RNA-Seq data also identified 5 different clusters which well correlated with the enhancer subtypes (**Fig. 3D** and **Extended Data Fig. 4B**) suggesting likely causative roles for enhancers in driving specific transcriptomic phenotypes. Consistently, the Variation of Information (VI) analysis showed that the enhancer patterns correlated best to the cancer type and mRNA expression patterns (**Fig. 3E**). To determine if these clusters were also present in tumor cells, and were not a result of artifact of cell culture, we overlapped EpiC-specific enhancer peaks with the ATAC-seq peaks from the TCGA tumors as well as H3K27ac peaks from 6 different tumor types (generated in house^26,28^ or publicly available^25,27^) **(Fig. 3F-3J** and **Table S6)**. Here we noted striking similarities in tissue-type composition. Similar to EpiC1 composition of mostly melanoma cell lines (**Fig. 3C**), the top raked tumors based on relative EpiC1 enrichment were all melanoma tumors **(Fig. 3F)**. Similarly, EpiC2 features were enriched in BRCA and OVCA tumors **(Fig. 3G).** Importantly, we also noted prostate adenocarcinomas to be enriched in this group suggesting EpiC2 to consist of hormone-driven cancers. EpiC3 features were enriched in COAD and STAD suggesting specificity of this cluster to GI cancers **(Fig. 3H)**. Consistent with pan-cancer nature of EpiC4 cell lines, tumors enriched in EpiC4 enhancers were pathologically defined to arise from multiple different tissues **(Fig. 3I)**. Finally, EpiC5 features were enriched in multiple gliomas as well as thyroid gland tumors **(Fig. 3J).**

**Fig. 3:**
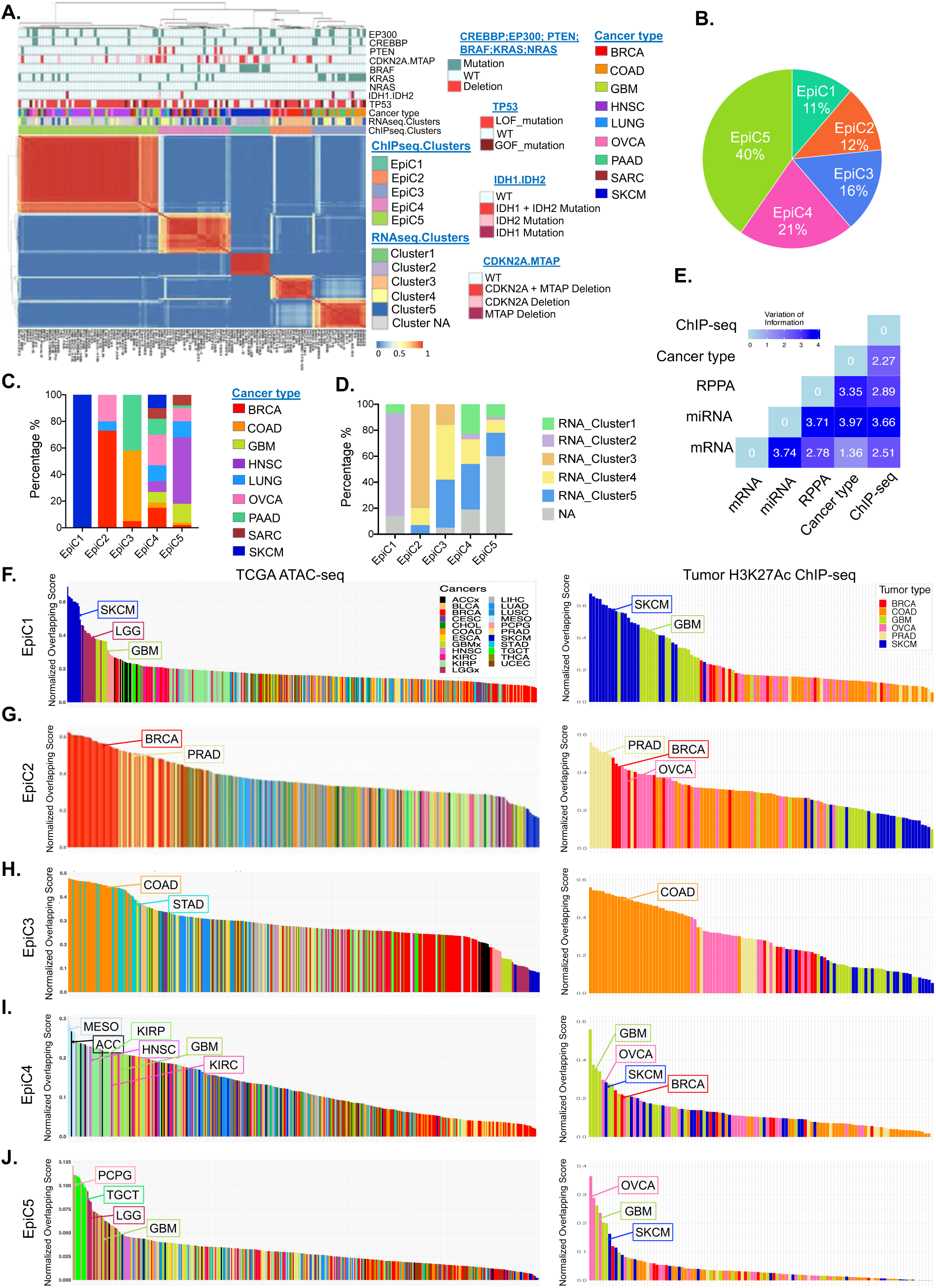
H3K27ac-defined enhancers classify cancer cell lines into distinct clusters. A. Heatmap of NMF-based clustering for H3K27ac enhancer data from cancer cell lines that defines the EpiC1-5 clusters. Top panel shows EpiC annotations and overlaps with RNA-seq NMF subtype (from S5B), cancer type and key oncogene mutation/CNVs. The cell line names are shown on the bottom. Each row of the heatmap represents an enhancer region; each column represents a cell line. Colors represents the H3K27ac ChIP-seq signal intensity. B. Pie chart showing the percent of cells in each EpiC cluster to total cell lines. **C-D**. Bar plot showing distribution of cancer subtypes (**C**) and RNA clusters (**D**) of the cell lines in each EpiC cluster. Colors represents the cancer type and RNA cluster type shown to the right of the plot. **E**. Variation of information (vi) analysis of clustering schemes derived by using various data types from CCLE. **F-J**. Bar plot showing the normalized overlapping score of EpiC1-5_specific enhancers with accessible chromatin regions (from ATAC-seq data) in the TCGA samples. Colors represents the cancer types shown above the plot. **K**. Heatmap showing the normalized overlapping score of EpiC1-5_specific enhancers with tumor H3K27Ac enhancer data.

### Molecular features of EpiC clusters

To further gain biological insights into the enhancer-based classification of cancer cell lines, we performed integrative analyses using existing omic datasets including transcriptomic (RNA-Seq), somatic mutations, proteomic (RPPA data), functional dependency (CRISPR screening through AVANA) and drug response^21,33^. Pathway analysis of EpiC-specific enhancer target genes, derived based on a previously defined enhancer-gene relationship as a function of three-dimensional contacts^34^, showed enrichment of specific pathways in each cluster: focal adhesion and IGF1/AKT signaling in EpiC1, ATM and AHR signaling in EpiC2, ERRB2 and EDA signaling in EpiC3, PI3K-AKT and focal adhesion in EpiC4, and PDGF pathway and Notch signaling in EpiC5 (**Fig. 4A**). Interestingly, those TCGA tumors have high overlap with EpiC-specific enhancers in each cluster showed enrichment of cancer type specific pathways in EpiC1-3, consistent PI3K-AKT pathway enrichment in EpiC4 and Notch signaling in EpiC5 (**Extended Data Fig. 4C-4G).**

**Fig. 4:**
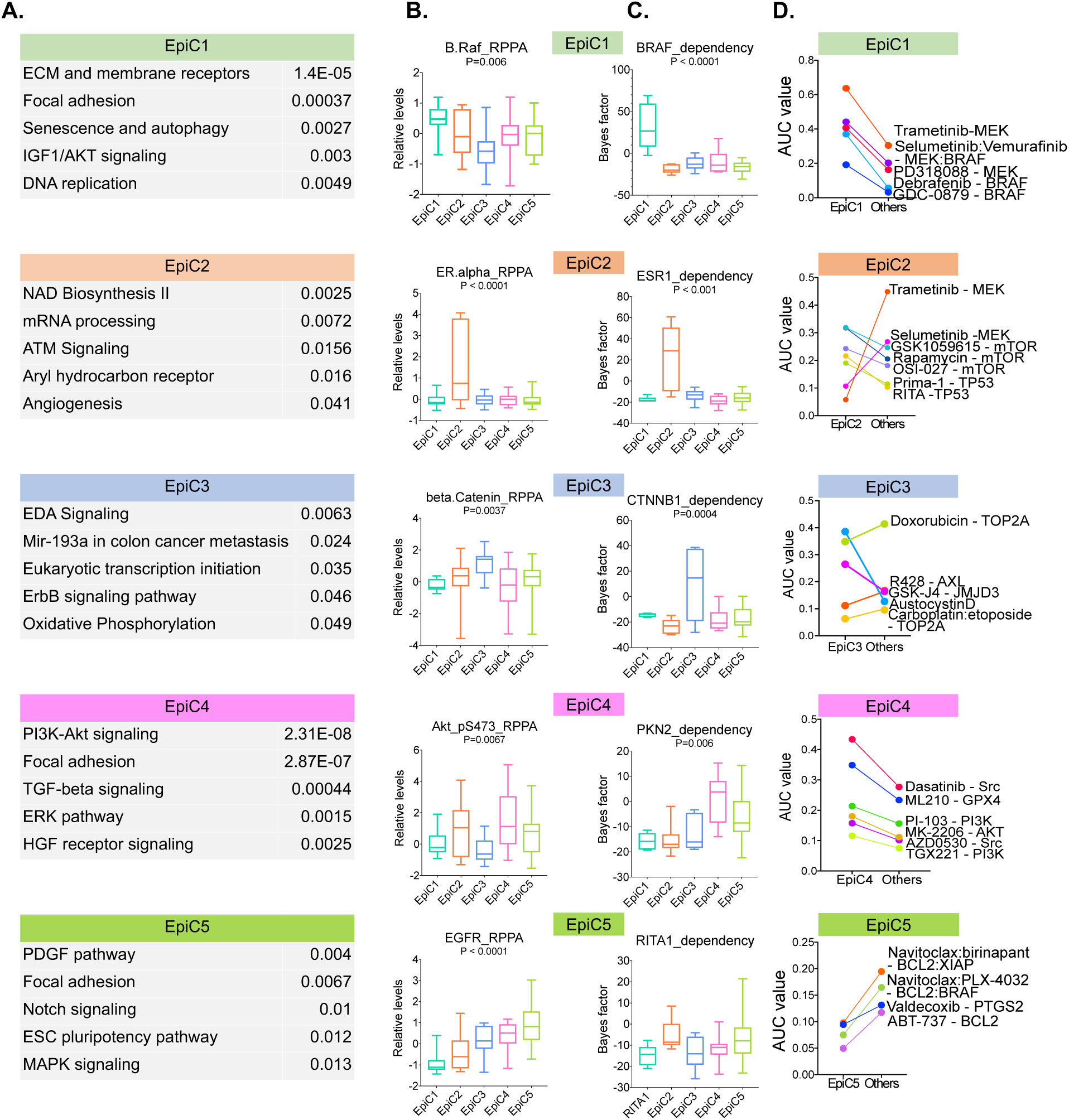
EpiC clusters display activation of specific signal pathways and unique responses to targeted therapies. **A.** Plot showing selected pathways from top 10 significantly enriched pathways (Enrichr) for EpiC-specific enhancers. **B.** Box plot showing relative levels for EpiC-specific proteins from CCLE RPPA dataset. Colors represents the EpiC clusters shown above the plot. P-values represent one-way ANOVA comparison between 5 EpiC clusters. **C.** Box plot showing Bayes factor for EpiC-specific dependency genes from AVANA dataset. Colors represents the EpiC clusters shown above the plot. P-values represent one-way ANOVA comparison between 5 EpiC clusters. **D.** Dot plots showing the average AUC values for the compounds showing significantly differential values in each EpiC cluster versus others. Colors represents the compounds shown above the plot. In (**B-C**), the bottom and the top rectangles indicate the first quartile (Q1) and third quartile (Q3), respectively. The horizontal lines in the middle signify the median (Q2), and the vertical lines that extend from the top and the bottom of the plot indicate the maximum and minimum values, respectively.

We did not observe correlations of EpiC clusters with somatic mutations in specific epigenetic regulators or bona fide oncogene and tumor suppressors such as TP53, CBP, EP300, IDH1, IDH2, KMT2C, KMT2D, PTEN, NRAS, SMARCA4, PBRM1, ARID1A, or ARID2 (**Fig. 3A** and **Table S4)**. Consistent with abundance of melanoma cell lines in EpiC1, we observed enrichment of BRAF mutations (**Fig. 3A)**, higher BRAF protein levels (**Fig. 4B** and **Extended Data Fig. 5A,** total and pS445), higher MAPK and ERK phosphorylation (**Extended Data Fig. 5A**) and higher dependency on BRAF, MAPK1 and TFAP2A in associated cells (**Fig. 4C** and **Extended Data Fig. 5B**). EpiC2, which is primarily composed of in breast cancer and ovarian cancer cell lines, showed higher protein levels of ER◻, HER2 and GATA3 and dependency on ER◻, GATA3 and ZNF217^35,36^, but no significant correlation of any mutations (**Fig. 3A, 4B-4C** and **Extended Data Fig. 5A-5B**). Similarly, consistent with prevalence of pancreatic and colorectal cancer lines in EpiC3, we observed higher levels of ◻-catenin and dependency on ◻-catenin and TCF7L2 (**Fig. 4B-4C** and **Extended Data Fig. 5A-5B**) as well as enrichment of KRAS mutations and dependency on KRAS in associated cell lines (**Fig. 3A** and **Extended Data Fig. 5B**). Consistent with enrichment of PI3K-AKT signaling pathway in EpiC4, we noted that 5 out of 6 cell lines harboring PTEN deletion were classified in this cluster (n = 26) (**Fig. 3A**). However, rest 21 cell lines did not show overt genomic features related to PI3K signaling thus suggesting potential activation of this pathway through enhancer activation. Consistently, proteomic data (RPPA) showed that the EpiC4 cell lines harbored higher levels of phosphorylated AKT (at S473 and T308) in comparison to other EpiC clusters (**Fig. 4B**). We also noted activation of PKCa in EpiC4 (**Extended Data Fig. 5A)** and dependency of EpiC4 lines on PKN2 (**Fig. 4C)**; both of these kinases are known to regulate AKT signaling^37,38,39^ (**Fig. 4C**). Finally, EpiC5 showed increased levels of EGFR as HNSCC and GBM cells lines were prevalent in this cluster (**Fig. 4B**). Consistent with deregulation of Notch signaling in HNSCC^40^, we noted dependency on a Notch regulator RITA1^41^ in EpiC5 (**Fig. 4C**).

Overlap of drug response data showed dependency of EpiC1 on MEK inhibitors whereas EpiC2 showed resistance to these inhibitors (**Fig. 4D**). However, EpiC2 showed sensitivity to mTOR inhibitors (**Fig. 4D**). Interestingly, EpiC3 showed dependency on JMJD3 inhibitors and resistance to topoisomerase inhibitors (**Fig. 4D**). Consistent with activation of PI3K pathway, we observed sensitivity to the AKT and PI3K inhibitors in EpiC4 cell lines (**Fig. 4D**). Finally, EpiC5 cell lines appear to be resistant to BCL2 inhibitors and its combinations (**Fig. 4D**).

### Dependence of EpiC Clusters on Specific Class of TFs

Since super-enhancers (SEs) upregulate the expression of important oncogenes in cancers^42^, we identified SEs unique to each EpiC cluster and determined their target genes (**Extended Data Fig. 6A-6E** and **Table S7**). Enhancers provide platforms for the binding of key TFs that act as drivers of the associated phenotypes. Hence, we defined “core TFs” that likely act as key regulators of the EpiC clusters by employing a previously published logic^43^. Here, we identified and integrated datasets including; 1) TFs defined by motif enrichment analysis (**Fig. 5A** and **Table S7**); 2) gene expression data for each EpiC (**Fig. 5A** and **Table S7**); 3) TFs marked by SEs for determination of self-regulation (**Fig. 5A** and **Table S7**); and 4) dependency data from the ACHILLES project of Broad/Novartis^33^ that was used to identify genes essential in all cell lines within each EpiC cluster (**Fig. 5A** and **Table S7**). This analysis identified several core TFs in each EpiC cluster (**Fig. 5A** and **Table S7**), including SOX10 and MITF in EpiC1, SPDEF and FOXA1 in EpiC2, CDX2 in EpiC3, ZEB1 in EpiC4 and ZBTB7A in EpiC5 (**Fig. 5B-5C** and **Extended Data Fig. 6B-6C**). Consistently, we observed the H3K27ac-marked enhancer peaks in the same regions in corresponding tumor samples (**Extended Data Fig. 6F**). Importantly, the identified TFs associated with these SE displayed “high” dependency (Bayes factor) and cluster-specific gene expression (**Fig. 5D**). We also observed that core TFs of the three tissue-type based clusters (EpiC1-3) have high expression in matched TCGA tumor types (**Extended Data Fig. 7A-7C**) whereas EpiC4 core TF ZEB1 is highly expressed in multiple TCGA tumors (**Extended Data Fig. 7D).** In addition, core TFs of these three clusters have significant dependencies in major cancer types compared with all other cancer types (**Extended Data Fig. 7E-7G**). Taken together, we identify key elements of core transcriptional circuitry that could be potential targets for EpiC-clusters.

**Fig. 5:**
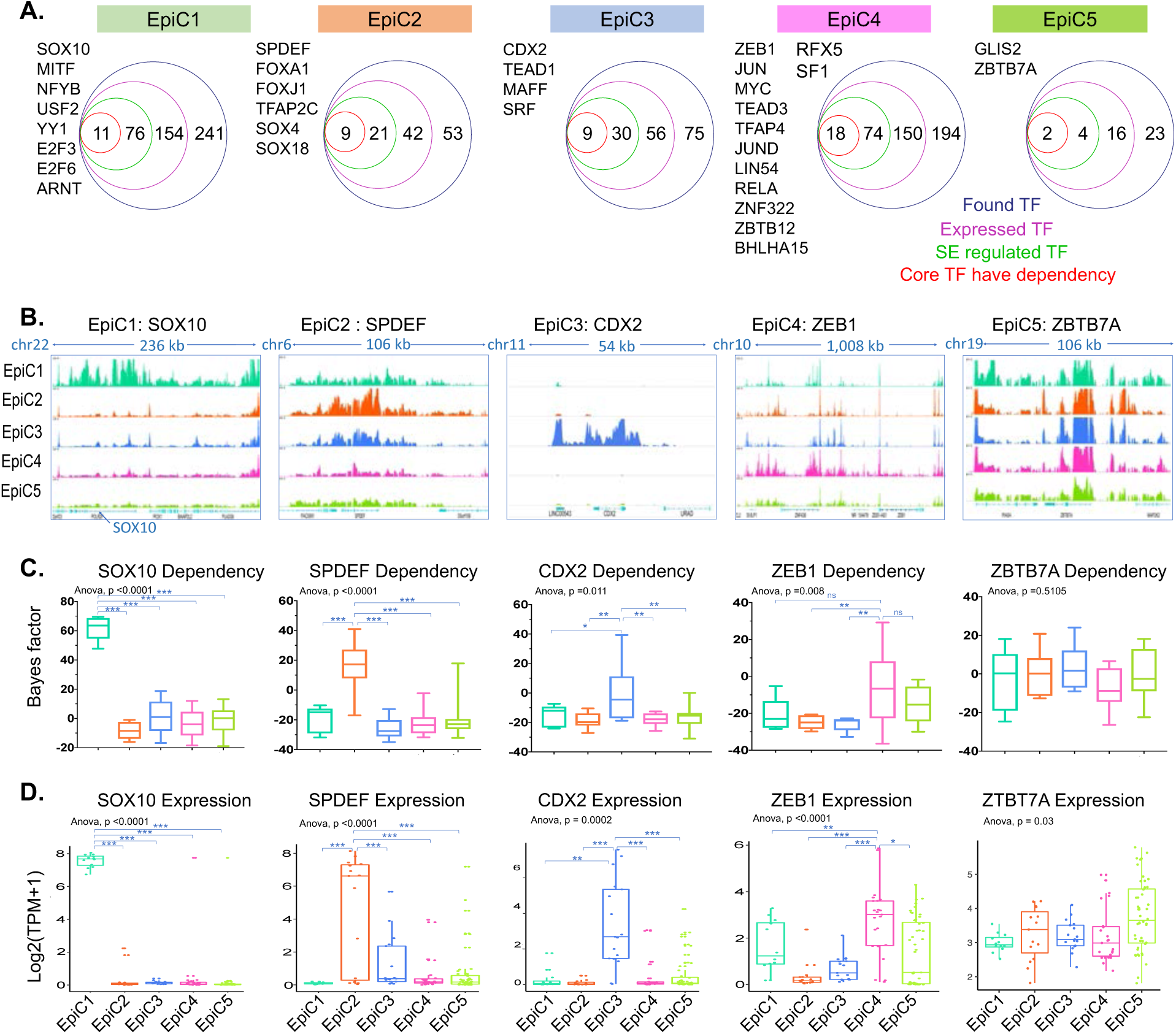
Super-enhancers regulates distinct core TF activities in each cluster. **A.** Core TF lists, which were transcribed, regulated by super-enhancers, and dependent in cell lines in the EpiC clusters. **B.** Genome browser view of composite ChIP-Seq tracks for H3K27ac surrounding representative EpiC-specific core TF genes: SOX10 for EpiC1, SPDEF for EpiC2, CDX2 for EpiC3, ZEB1 for EpiC4, and ZTBT7A for EpiC5. **C-D**. Box plot showing Bayes factor for dependency from AVANA dataset (**C**) and gene expression level (**D**) for EpiC-specific core TFs shown above. Colors represents the EpiC clusters shown above the plot. P-values represent one-way ANOVA comparison between 5 EpiC clusters. Unpaired t-test p-values were calculated for specific EpiC vs others. Asterisk denotes *p < 0.05, ** p < 0.001, *** p < 0.0001. The bottom and the top rectangles indicate the first quartile (Q1) and third quartile (Q3), respectively. The horizontal lines in the middle signify the median (Q2), and the vertical lines that extend from the top and the bottom of the plot indicate the maximum and minimum values, respectively.

### EpiC4 tumors display poor survival

Investigation of clinical data for the TCGA tumors that showed enrichment for various EpiC enhancer signatures showed that the tumors harboring enrichment of EpiC4 signature had significantly poor survival compared with those that did not (**Fig. 6A**). In contrast, the EpiC2-related TCGA tumors showed better survival, whereas tumors corresponding to other clusters showed no significant differences (**Extended Data Fig. 8A-8D**). Since EpiC4 is a novel pan-cancer enhancer-based subtype with poor clinical outcomes, we further analyzed the molecular pathways in these cells. Consistent with activation of PI3K pathway, we observed higher area under curve (AUC) values of multiple PI3K inhibitors (CAL-101, TGX221, PI-103, and GSK2636771) in EpiC4 cell lines compared to those in other clusters (**Fig. 6B**). Similarly, consistent with higher AKT phosphorylation, we had observed higher AUC values for AKT inhibitor, MK-2206, in EpiC4 lines (**Fig. 6B**). The tumor types that had higher overlaps in EpiC4 included ACC, MESO, LGG, and GBM (**Fig. 3I**), which harbored high PI3K-AKT/mTOR scores^44^. Deeper look into the regulatory networks enriched in EpiC4 suggested activation of SRC kinase protein network (**Fig. 6C**) which was is likely a driver event as suggested by the dependency on various members of this network (**Fig. 6D**) as well as sensitivity of EpiC cell lines to SRC-specific inhibitors, Dasatinib and AZD-0530 (**Fig. 6E**). Furthermore, RPPA data suggested activation of E-cadherin, Caveolin and Fibronectin that are known to merge into SRC kinase activation in aggressive cancers driving invasion and motility of cancer cells (**Fig. 6F**) ^45,46,47,48,49,50^. Thus, we propose that EpiC4, an aggressive pan-cancer enhancer-based subtype of cancers, is driven by a distinct Fibronectin-CAV1-SRC-PI3K-AKT axis (**Fig. 6G**). We also noted activation of specific networks in different EpiC clusters including AHR-driven network in EpiC1 (**Extended Data Fig. 8E-8F**), GRHL2-driven network in EpiC2 (**Extended Data Fig. 8G-8H**), KLF5-driven network in EpiC3 (**Extended Data Fig. 8I-8J**) and RAC-1 driven network in EpiC5 (**Extended Data Fig. 8K-8L**). Overall, distinct enhancer-based subgroups of cancer cells show activation of specific transcriptional programs and responsiveness to different pharmacological agents which can be exploited for therapeutics in future based on their unique enhancer signature (**Fig. 7**).

**Fig. 6:**
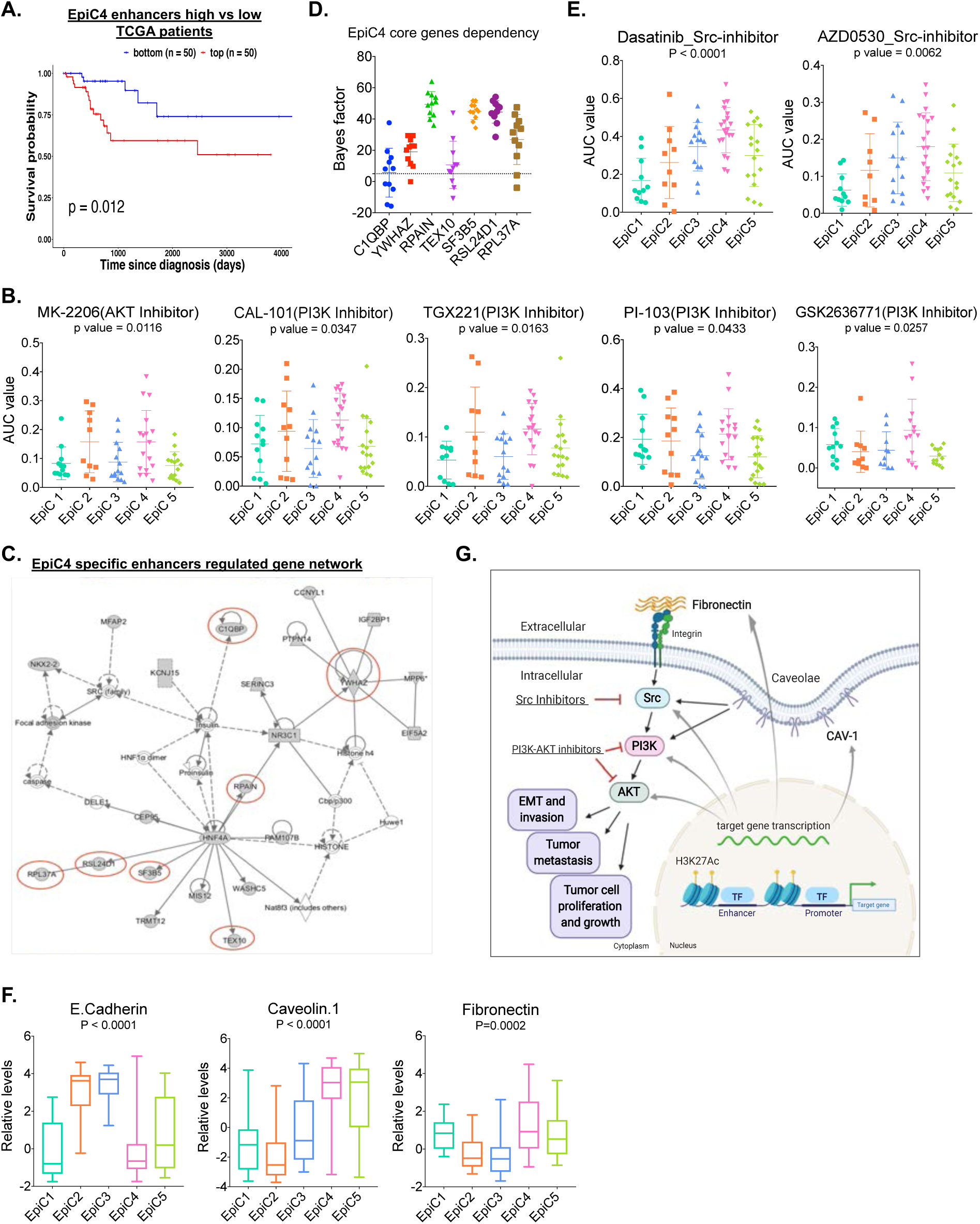
EpiC4 is a novel pan-cancer enhancer cluster with mesenchymal signature. **A.** Kaplan-Meier analysis showing overall survival of patients (whose samples were profiles by the TCGA ATAC-Seq) with top 50 and bottom 50 normalized overlapping scores as shown in Fig. 3F. p-value is derived from log rank test. **B.** Dot plots showing the AUC values for PI3K and AKT inhibitors in each EpiC cluster. Each dot represents a given sample. Colors represents the EpiC clusters shown above the plot. P-values represent one-way ANOVA comparison between 5 EpiC clusters. **C.** IPA gene network analysis based on the EpiC4 specific enhancer annotated genes. Red circled genes have dependency in EpiC4 cell lines. **D.** Dot plot showing Bayes factor for the genes from the panel C network for EpiC4 cell lines. **E.** Dot plots showing the AUC values for Src inhibitors in each EpiC cluster. Each dot represents a given sample. Colors represents the EpiC clusters shown above the plot. P-values represent one-way ANOVA comparison between 5 EpiC clusters. **F.** Box plot showing the levels of E-cadherin, caveolin and fibronectin from RPPA data in EpiC clusters. Colors represents the EpiC clusters shown above the plot. P-values represent one-way ANOVA comparison between 5 EpiC clusters. The bottom and the top rectangles indicate the first quartile (Q1) and third quartile (Q3), respectively. The horizontal lines in the middle signify the median (Q2), and the vertical lines that extend from the top and the bottom of the plot indicate the maximum and minimum values, respectively. **G.** Schematic illustrating cluster specific enhancers regulated Fibronectin-CAV1-SRC-PI3K-AKT axis in EpiC4.

**Fig. 7:**
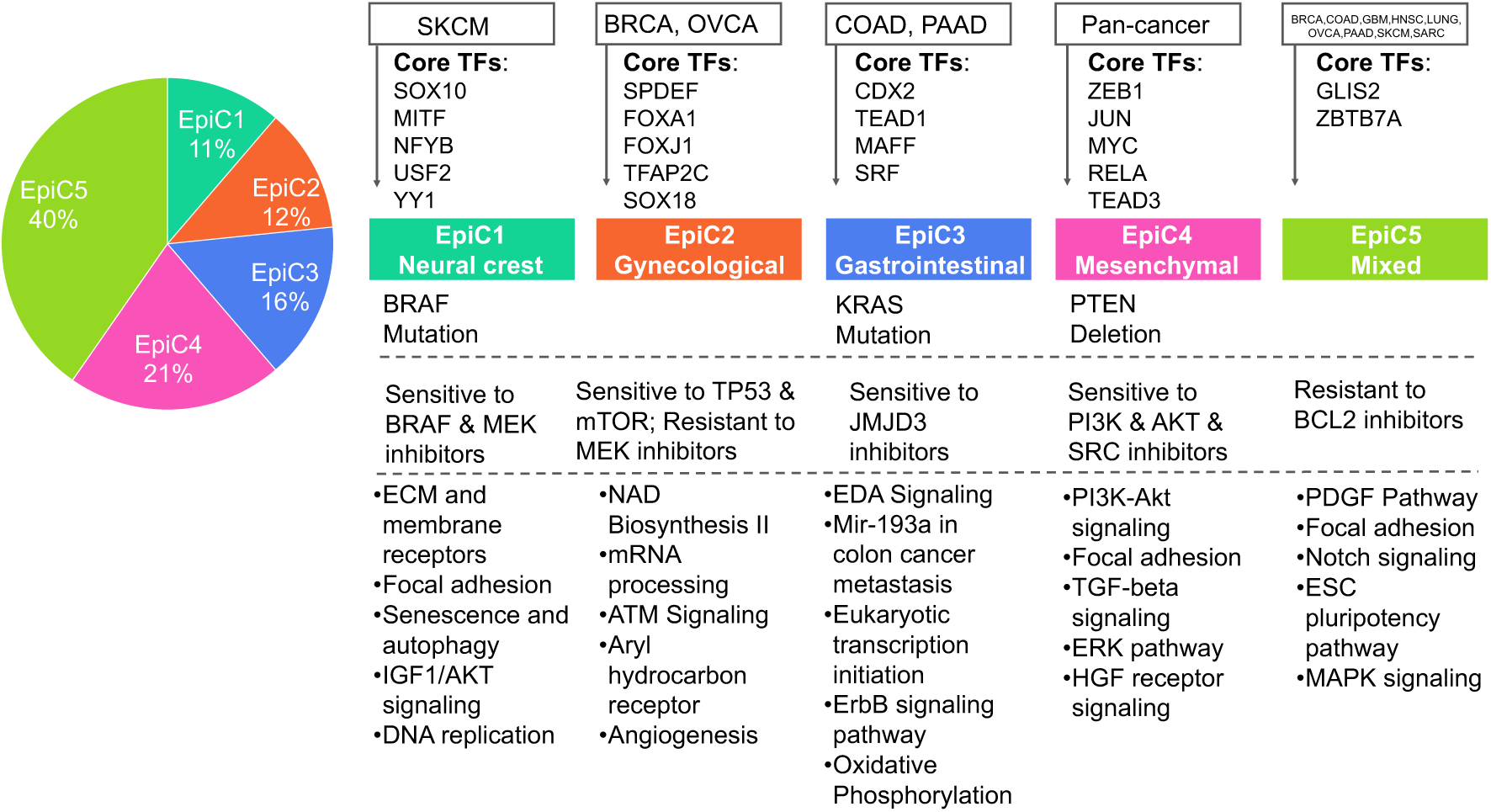
Enhancers of cancer cell lines identifies specific clusters and targetable dependencies. Schematic illustrating the summary of findings for the 5 EpiC-clusters identified in this study.

## DISCUSSION

Our data and analyses provide initial evidence for several novel phenomena that need to be studied in-depth in the future. Firstly, we define a novel enhancer based subtype, EpiC4, for which corresponding patients show poor survival and unique features that can be exploited to devise therapeutic strategies in the future. Secondly, our results identify the concept of core TFs for enhancer-based subgroups that could be exploited in future with novel therapeutic strategies such as PROTAC^51^. Thirdly, we provide initial evidence that specific non-coding epigenetic elements, such as enhancers, could potentially serve as therapeutic targets for overcoming resistance to FDA-approved agents in specific cancers. Here, advances in RNA therapeutics could be exploited to target enhancer RNAs.

We presented comprehensive profiles of chromatin modification maps from a large number of widely used cancer cell lines with associated multi-omics datasets (mutation, copy number, gene expression, methylation, RPPA, ACHILLES and drug response data). While cancer cell lines have been passaged extensively, yet they harbor most of the same genetic changes found in patients’ tumors^22^; therefore, they have been used extensively as good models for dissecting relationships between genetic aberrations and drug responses. However, it is not known whether the epigenetic landscape of these heavily passaged cell lines in various culture media retain the epigenetic features from the human tumors. We demonstrate that H3K27ac peaks between cell lines and human tumors show 53-77% overlaps. Furthermore, H3K7ac-peaks overlap at ∼60% frequency with open chromatin regions defined by ATAC-Seq in the TCGA tumors. Even though H3K27ac and ATAC-Seq are two different methods for identifying enhancers, and tumors harbor multiple different cell types, this significant overlap between our data and that of the TCGA ATAC-Seq study suggests that these elements are truly active in our dataset, and these cancer cell lines remain a good model for studying cancer-specific chromatin aberrations.

Using combinatorial chromatin state analysis, we demonstrated that enhancer deregulation is a fundamental epigenetic feature of cancer cells. In-depth investigation of enhancer patterns identified five epigenetic subtypes (EpiC1-5), in which we identified targetable core TFs that play a critical role in the biology of each specific EpiC cluster. By integrating these enhancer peaks with drug response data, we identified novel enhancer regulated gene dependencies of specific cancer types that are responsive to pharmacologic inhibition. Most notably, we identified a new epigenetic cluster (coined “EpiC4”) that consists of cell lines from all 9 tumor types, suggesting that a subgroup of each cancer type likely harbors a predominant component of enhancer aberrations that are more distinguishable than ones defined by their tissue of origin. Based on our findings we propose that EpiC4 enhancer features are an underlying epigenetic feature of aggressive cancer types. EpiC4 cancer cells display mesenchymal characteristics that promotes invasion and metastasis, resulting in poor survival. This is supported by discovery of ZEB1 as a key driver for this subtype and upregulation of Cav1-SRC1-PI3K-AKT pathway which promote epithelial-to-mesenchymal transition (EMT)^48,46,52,53,54^.

Pluripotency factors such as OCT4, SOX2 and NANOG have pioneering ability to access chromatin and initiate marking of active chromatin around potential binding sites for key TFs required for maintaining stem-cell renewal or differentiation, and therefore are called ‘core TFs’. These TFs also self-regulate their own expression through generation of SEs in the vicinity of their gene promoters. In this study, the identification of distinct EpiC clusters provides insight regarding the functional role of core TFs and their interplay with enhancer and gene expression patterns in multiple cancer types. While we identified anywhere from 2-18 TFs as core drivers in different EpiC clusters, whether these have true potential to set up enhancer patterns on their own or in cooperation with the other core TFs for that subgroup will need to be determined in an experimental setting. We believe that identified core TFs in each cluster provide a resource with which to explore the core TF directed therapeutic interventions as they show specific dependency in that EpiC group. For example, SOX10 could be an attractive target for melanoma, given the dependence of the neural crest and melanocytic differentiation on this important pioneer TF^55^. Recent advances in PROTAC technology^51^ that allow targeting of TFs for degradation by the proteasome may be exploited to devise new therapeutic approaches based on our identification of core TFs specific for EpiC subgroups.

Overall, our results represent a comprehensive attempt to describe the chromatin landscape in cellular models of nine solid tumor types using representative cancer cell lines. The data resource and analyses described here demonstrate that knowledge of chromatin state landscapes can be used to dissect the epigenetic similarities and differences between multiple cancer lines and to provide unique information that may inform future precision therapies.

## ACKNOWLEDGMENTS

We thank Kadir C. Akdemir for sharing and helping with processing the HCC1954 Hi-C data. We thank Curtis Gumbs, SMF Core and ncRNA and Sequencing Core at The University of Texas MD Anderson Cancer Center for sequencing support. The work described in this article was supported by grants from the National Institutes of Health (CA160578, CA226269 to K.R.; CEP awards from Bladder, Lung, Brain, Head&Neck SPOREs to K. R.; CA016672 to SMF Core), Department of Defense (NF190074 to K.R. and NF190050, NF160026 to K.E.T.), Center for Cancer Epigenetics at MD Anderson (K.R.), CPRIT (Cancer Prevention & Research Institute of Texas; RP170407 and RP200390 to K.R.), Kadoorie Reseach Funds to J.M. and MD Anderson Cancer Center (start-up funds to K.R.). S. Srinivasan was supported by the CPRIT Research Training Grant (RP170067). The histology work was performed at the Histopathology Core Lab at the UT MD Anderson Cancer Center supported by the NIH National Cancer Institute (P30CA016672).

## AUTHOR CONTRIBUTIONS

M.M. conceptualized and designed the study, planned and conducted the experiments, analyzed data, prepared figures, and wrote the manuscript. C.J.T., Z.L., E.O. and V.K. generated ChIP-Seq data for a subset of cell lines. E.A. performed integrative analyses of enhancer data with drug response, ATAC-Seq and other omic datasets. A.T.R. conceived and performed NMF clustering analysis for enhancer and RNA-Seq data, generated the ChromXplorer tool, and helped with chromatin state analyses. J.S., S. Srinivasan, M.T., S.B.A., J.E.M.L. and K.T. helped with bioinformatics analysis. A.K.S., N.S. and M.C. provided technical help. S. Sarkar contributed to study design. K.E.T., F. M. J., C.R. P., C.B. and J.M. provided reagents and support for fellows. K.R. conceived, designed and conceptualized the study, evaluated data, prepared the figures, and wrote the manuscript.

## METHODS

### Cell Lines

Cell lines profiled by CCLE/Sanger and MDACC (Table S1) were grown according to vendor recommendations. All cell lines were verified by STR profiling and checked for mycoplasma intermittently using a PCR based MycoAlert Mycoplasma detection kit (Sigma).

### Chromatin Immunoprecipitation

Chromatin immunoprecipitation was performed as previously described^1^ using 3 million cells per sample and 5mg of antibody per ChIP experiment for H3K4me1 (Abcam ab8895), H3K27ac (Abcam ab4729), H3K4me3 (Abcam ab8580), H3K9me3 (Abcam ab8898), and H3K79me2 (Abcam ab3594) or 8mg of antibody per ChIP experiment for H3K27me3 (Abcam ab6002). Enriched DNA was quantified using Qubit (Thermo Fisher Scientific), and ChIP libraries were amplified and barcoded with use of the NEBNext® Ultra™ II DNA library preparation kit (New England Biolabs) according to the manufacturer’s recommendations. After library amplification, DNA fragments were size-selected (200 - 600 bp) using AMPure XP beads (Beckman Coulter) and assessed using high sensitivity D1000 screentape on the Bioanalyzer (Agilent Technologies). Libraries were sequenced at the Advanced Technology Genomics Core (MD Anderson) or at the Cancer Genomics Laboratory using Hi-Seq 2000 or HiSeq 4000 (Illumina) instruments in the 36bp or 50bp single-end read format.

### ChIP-seq Data Processing and Alignment

ChIP-seq data were quality controlled and processed by pyflow-ChIPseq^2^, a snakemake^3^ based ChIPseq pipeline. Briefly, a list of fastq files for the same mark were merged first. After merging, FastQC was used to check overall sequence data quality. Bowtie^4^ was used to align to the hg19 human reference genome using “--chunkmbs 320 -m 1 --best -p 5” options. SAMtools was used to sort and isolate uniquely mapped reads using “samtools sort -m 2G -@ 5” options. Duplicated reads were removed by SAMBLASTER, and only uniquely mapped reads were retained. This resulted in the final aligned, de-duplicated BAM file that was used in all downstream analyses.

### ChIP-seq Data Quality Control

To avoid artificial differences in signal strength due to differences in sequencing depth, all consolidated histone mark data sets were uniformly subsampled to a maximum depth of 15 million reads. These uniformly subsampled data sets were then used for all further processing steps (peak calling, signal coverage tracks, chromatin states). Phantom peak quality checks used to generate quality metrics: Normalized strand cross-correlation (NSC) and relative strand cross-correlation (RSC); NSC and RSC metrics use cross-correlation of stranded read density profiles to measure enrichment independently of peak calling. RPKM normalized bigwigs were generated by deeptools^5^ and checked quality of each dataset across the genome on the Integrative Genomics Viewer ^6^.

### Peak calling

For narrow peaks, the macs1.4^7^ peak caller was used to compare ChIP-seq signal to a corresponding input sequenced control to identify narrow regions of enrichment (peaks) that pass a p-value threshold of 1e^-5^. For broad domains, the MACSv2.0.10 peak caller was used with a -- broad-cutoff p-value of 1e^-5^.

### Chromatin State Analysis

ChromHMM^8^, which is based on a multivariate Hidden Markov Model, was used with default parameters to derive genome-wide chromatin state maps for all cell types. For each consolidated ChIP-seq data set, read counts were computed in non-overlapping 1000-bp bins across the entire genome. Each bin was discretized into two levels, 1 indicating enrichment and 0 indicating no enrichment. The binarization was performed by comparing ChIP-seq read counts to corresponding input control read counts within each bin and using a Poisson p-value threshold of 1e^-4^. Chromatin state models were learned jointly on all data for all 6 histone marks (H3K4me1, H3K4me3, H3K27ac, H3K79me2, H3K9me3, and H3K27me3) from 80 highest quality cancer cell lines and a model with 15 states was chosen for detailed analysis since it captured all the key interactions between the chromatin marks. The trained model was then used to compute the posterior probability of each state for each genomic bin in each reference epigenome. The regions were labelled using the state with the maximum posterior probability. To determine the chromatin state differences between different groups, we used a two-step process. First, using the segmentation calls from the ChromHMM output, the entire genome was divided into non-overlapping windows of 10 kb. We next counted the number of times a chromatin state was observed in each of the 10 kb windows and obtain a frequency matrix for each state in the ChromHMM model (E1-E15). In the second step, low variable genomic loci were removed from the frequency matrix, and significant differences between multiple groups of samples types were calculated by using a Kruskal-Wallis rank sum test with a p-value of 1e^-3^ for each state separately.

### t-SNE Analysis of Chromatin State and H3K27Ac Defined Enhancer Data

For unsupervised hierarchical clustering of each chromatin state for cancer cell lines, and for clustering cancer cell lines H3K27ac ChIP-seq data alone or with Roadmap normal samples together, we calculated the distance measure. The resulting distance matrix was used to perform the t-SNE analysis (t-Distributed Stochastic Neighbor Embedding, Rtsne package version 0.11) ^9^. The following non-default parameters were used: theta=0, dims = 2, max_iter=10000, perplexity = 30.

### Enhancer Diffbind analysis

The 125 of best quality H3K27ac ChIP-seq data sets were chosen for quantitative comparison using the Diffbind^10^ package. Only peaks present in at least 2 or more samples were retained. The peaks were then merged by Diffbind across samples, and a ChIP read counts minus Control read counts matrix was obtained. Peaks that overlapped with 2.5kb upstream and 2.5kb downstream of any known TSSs were removed.

### Super Enhancer Analysis

Super-enhancers (SEs) were defined by using the Rank Ordering of Super Enhancers (ROSE2) algorithm^11^. For all samples, the stitching distance was fixed at 12.5 kb to facilitate comparisons between samples. All other parameters used the default setting. The ChIPseeker package was used for defining the target genes of super-enhancers for subsequent analyses.

### Differential Enhancer Analysis

#### Overlap of Cancer enhancer peaks and Roadmap enhancer peaks

To identify variable enhancer domains enriched in either cancer cell lines or Roadmap normal tissues, we first defined the union of all H3K27AC enhancer peaks discovered across the cancer cell lines and normal tissues. Peaks that overlap with 2.5kb upstream and 2.5kb downstream of any known TSSs were then removed. Bedtools^12^ was used to identify overlap or unique peaks from the two datasets.

#### Comparison of enhancer peaks for cancer developmental stages

Comparisons of enhancer peaks were performed with BEDOPSv2.4.21^13^. BED files containing H3K27Ac ChIP-seq peaks for an individual sample were sorted with the sort-bed function. For downstream analyses, peaks were merged by developmental state (ESC, normal or Cancer) using the bedmap function to identify the shared peaks in at least 25% of the samples in a group. The unique and shared peaks within multiple groups were identified by Intervene^14^. The peaks were annotated with ChIPseeker R package^15^.

### Enhancer Data Analysis – Peak-to-Gene Linking Predictions

To identify putative causal links between enhancer peaks and gene expression, we used a reference-based approach. We downloaded previously published enhancer-target networks inferred by JEME from the ENCODE+Roadmap and FANTOM5 samples^16^ and overlapped our enhancer peaks with the reference enhancer to assign the enhancer to the refseq promoter and get the annotation. In addition, we also used the ChIPseeker package for annotation, using addFlankGeneInfo function for SEs.

### RNA-seq Data Analysis – Constructing a Counts Matrix and Normalization

Published RNA-seq data was downloaded from CCLE portal. To combine with MD Anderson home-generated RNA-seq data, we used the transcripts per million (TPM) normalized data for each cancer cell line, matching to the CCLE IDs with matched ChIP-seq. This was done first by length normalizing the read counts per gene by their respective exon lengths and then normalizing these values to a million within each sample. For gene names that were duplicated, we kept the genes that had the highest TPM variance. TCGA RNA-seq transcription comparison analysis within cancer types was performed on the UALCAN website^17^.

### Identification of Enhancer and Super Enhancer Associated Pathways

Differential enhancers and SEs associated genes in each cluster or in each cancer developmental stages were imported into the ClusterProfiler^18^ for pathway analysis, restricted to GO, KEGG, Hallmark and WiKi gene sets. The Enrichplot package ^19^ was used to generate dotplot and networks for genesets enriched with a false discovery rate (FDR) cut-off of < 0.05.

### Enhancer Region Identification at Positions of Cancer-Specific GWAS SNPs

Published genome-wide association studies (GWAS) associations were downloaded from the European Bioinformatics Website (https://www.ebi.ac.uk/gwas/docs/file-downloads). All GWAS associations for our ChIP-seq data matched cancer types were separated and then overlapped with each cancer type merged enhancer peaks to identify those GWAS SNPs located on an enhancer region of each cancer type.

### Variation of Information Analysis

mRNA, miRNA, and RPPA were downloaded through Broad Institute’s data portal (https://portals.broadinstitute.org/ccle/data). After preprocessing and normalization steps, nmf clustering was applied for all datasets to be consistent. The optimal number of clusters was searched between 2 to 15 for all datasets. Variation of Information is a distance measurement, using the summation of conditional entropies. The intuition for a conditional entropy is how much additional information is needed to predict a set by a given set. So, if VI is low when we compare two clustering results, it means they are similar. We used the mcclust R package to get all VI scores.

### Unsupervised Hierarchical Clustering Analysis of Variant Enhancer Loci in Cancer Cell Lines

A matrix of the normalized H3K27AC minus control read counts was obtained in Diffbind based upon the consensus typical enhancers (TEs) identified. In the case of unsupervised hierarchical clustering between 124 cancer cell lines, the top 10,000 variant enhancer loci were retained by using the *FSbyMAD* function in CancerSubtypes package in R. These enhancers were used to perform consensus nonnegative matrix factorization (NMF) using NMF package in R ^20^. Based on the dispersion, cophenetic and silhouette scores, k, the optimal cluster number of 5 was determined in both ChIP-seq and RNA-seq datasets.

### Enhancer Region Overlap with Tumor H3K27Ac Enhancer Peaks

Ovarian cancer and triple negative breast cancer tumor H3K27Ac ChIP-seq data were downloaded from the Gene Expression Omnibus (GEO) repository (accession #GSE121103) and The European Bioinformatics Institute (EMBL-EBI) repository (accession # PRJEB33558). Melanoma, colon cancer, prostate cancer and GBM H3K27Ac ChIP-seq data were generated in house and data have been deposited into the Gene Expression Omnibus (GEO) repository (accession #GSE 134043, #GSE136889). Five distinct epigenetic subgroups’ regions were overlapped with all profile’s H3K27Ac ChIP-seq peaks. Overlapping the matrix with the number of profiles within a cancer x 5 dimension was normalized by the different number of epigenetic subgroup-specific regions. Then we scaled peak overlapping scores so that 5 epigenetic subgroup-specific peak intersection scores totaled 1. For each subgroup, those normalized overlapping scores were ranked.

### Enhancer Region Overlap with TCGA ATAC-seq Peaks

Cancer type-specific normalized counts from “The chromatin accessibility landscape of primary human cancers” were used as the ATAC-Seq peaks source with 796 profiles and 23 different cancer types. Regions in that matrix are containing all of the reproducible peaks for the corresponding cancer. The appropriate threshold was defined by using the bigwig tracks for each cancer and peaks for each sample within a cancer were defined. Five distinct epigenetic subgroups’ regions were overlapped with all profile’s ATAC-Seq peaks. Overlapping the matrix with the number of profiles within a cancer x 5 dimension was normalized by the different number of epigenetic subgroup-specific regions. Then we scaled peak overlapping scores so that 5 epigenetic subgroup-specific peak intersection scores totaled 1. For each subgroup, those normalized overlapping scores were ranked.

### Enrichment of Motifs in Cluster-Specific Enhancer Peaks

To identify the motifs over-represented within each of the EpiC-specific enhancer peak sets, we used the HOMER motif database and the coordinates of EpiC-specific peak sets^21^.

### Identification of Core Transcriptional Regulatory Networks in Enhancer Clusters

Raw motif enrichment is insufficient to predict exactly which TF is mediating that activity. For this reason, we looked for the correlation of the motif’s –log_10_(p-value) and the expression of the transcription factor (log_2_(TPM + 1)) and identified those factors where |R| > 0.3 as potential mediators of the observed motif enrichment.

### Breast Cancer Known Subtype Calls

Breast cancer cell lines were classified into breast cancer gene expression-based subtypes based on previously published classifications ^22^.

### HiChIP Data Analysis

HiChIP experiments were performed as previously described by Mumbach et al.^23^, with minor modifications. HiChIP paired-end reads were aligned to the MboI digested hg19 genome using the HiC-Pro pipeline with default conditions. The default setting of HiC-Pro removes duplicate reads, assigns reads to MboI fragments, identifies valid interactions and generates hi-resolution interaction matrices. HiChIP for H3K27ac generated high-resolution contact maps containing ∼65 million valid interactions in HCT116 cells. Files for Juicebox visualization were generated using the HiC-Pro hicpro2juicebox.sh command based on the total valid interactions. Identification of H3K27ac mediated loops was performed with the hichipper/diffloop programs using the HiC-Pro output and ChIP-seq peaks from H3K27ac as anchor loci. Hichipper identifies intrachromosmal looping between anchor loci within 5kb-2MB and produces a per-loop FDR value from the loop proximity bias correction implemented by Mango. Using the mango output from hichipper, diffloop was used to filter significant loops (FDR < 0.01, width ≥ 5000, loop-count ≥ 2) and define enhancer-enhancer and enhancer-promoter interactions.

### Survival analysis

Clinical information for each sample was obtained by GDCquery_clinic from the TCGAbiolinks R package. Expression data were obtained from the NIH Genomic Data Commons. The survminer package was used for drawing the Kaplan-Meier plots and defining the optimal threshold (function surv). The outcome is overall survival censored at 10 years. P-values reported for the univariate model correspond to the logrank test.

### Quantification and statistical analysis

The two-tailed Student’s t-test was used to determine the statistical significance of two groups of data; One-way ANOVA was used to determine the statistical significance of three or more groups of data using GraphPad Prism. Data are presented as means ± standard error of the mean (SEM; error bars) of at least three independent experiments or three biological replicates. P-values of less than 0.05 were considered statistically significant. *, P <0.05; **, P <0.01; and ***, P <0.001 indicate statistically significant differences.

### Data and code availability

All ChIP-Seq dataset generated from cancer cell lines have been deposited into the Gene Expression Omnibus (GEO) repository (accession #GSE142751). All codes are available at https://gitlab.com/railab.

## EXTENDED DATA FIGURE LEGENDS

**Extended Data Fig.1:**
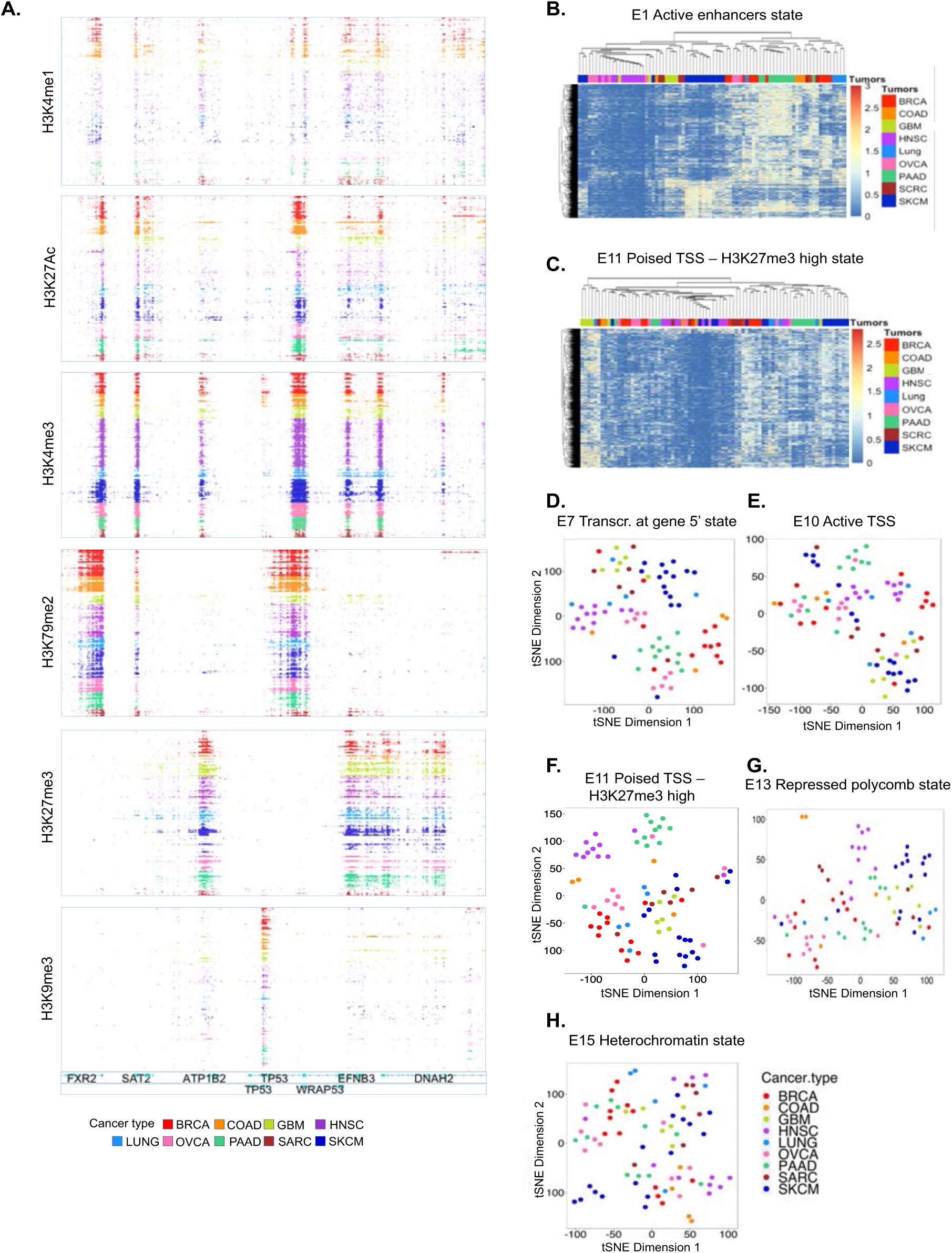
Chromatin state patterns of cancer cell lines. **A.** Genome browser views of ChIP-Seq tracks for 6 histone modification marks on the genomic loci surrounding TP53 gene to demonstrate the presence of active or repressive marks. Color represents the cancer types shown above the plot. **B-C.** Heatmap representation of the unsupervised clustering of the most variable “Active enhancer” state (**B**) regions or “Poised TSS – H3K27me3 high state” (**C**) across all cancer types. Each row represents a region, each column represents a cell line. Color represents the relative ChIP-seq signal for each region as a z-score. **D-H**. Unsupervised t-SNE on the top 50 principal components for the 10,000 most variable genic enhancers state (**D**), transcription at gene 5’ state (**E**), active TSS state (**F**), repressed polycomb state (**G**) and heterochromatin state (**H**) regions across all cancer types. Each dot represents a given sample. Colors represents the cancer type shown above the plot.

**Extended Data Fig.2:**
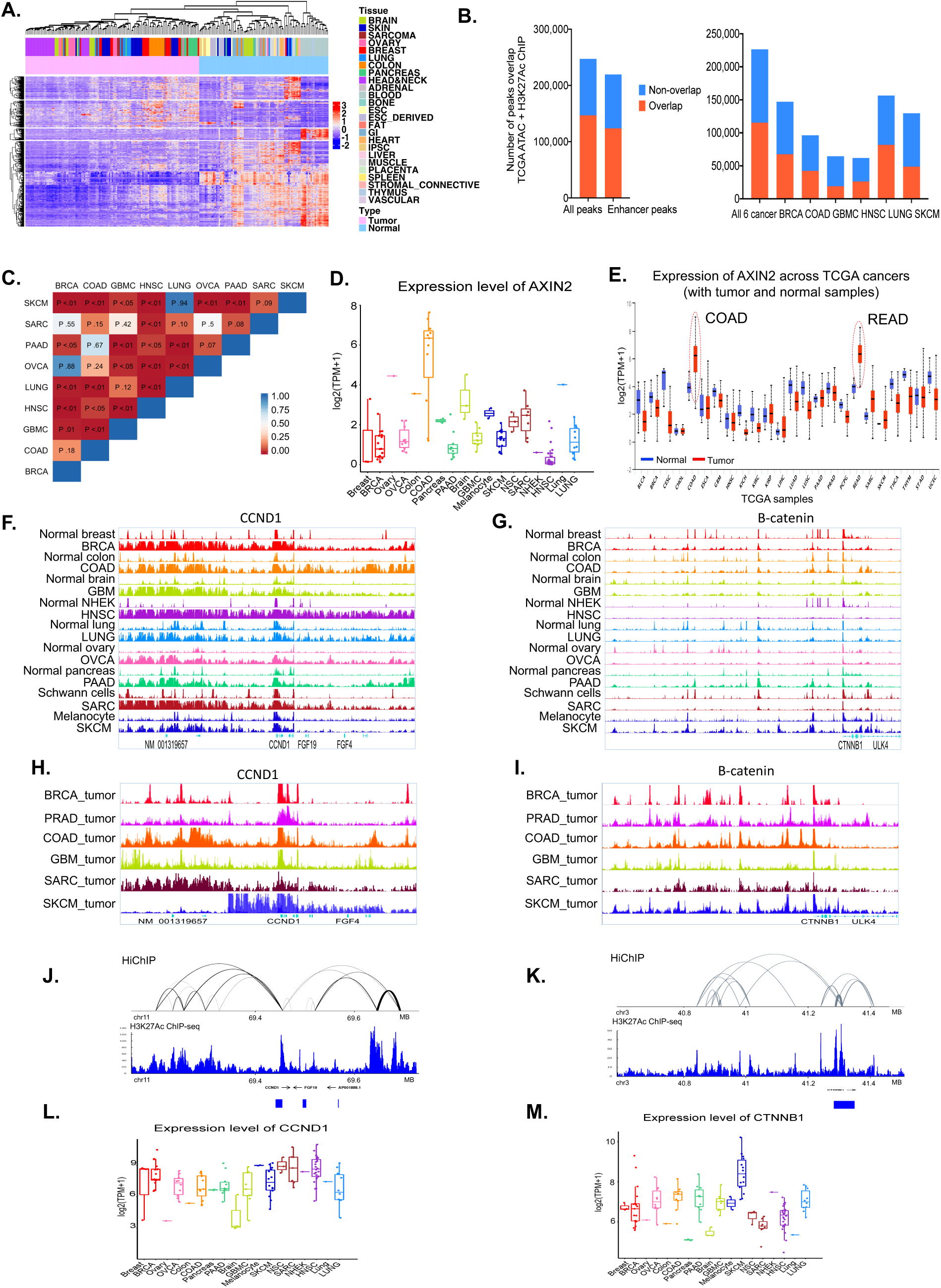
Enhancer deregulation in cancer cell lines. **A.** Heatmap representation of the unsupervised clustering of top 10,000 most variable enhancer peaks for all 124 cancer cell lines and 93 Roadmap normal samples. Each row represents an enhancer region; each column represents a cell line. Colors represents the H3K27Ac ChIP-seq signal RPKM value. **B.** Bar plot showing number of overlaps between H3K27Ac peaks from all cancer cell lines (labelled ‘All’) and those outside +/-2.5kb TSS (labelled ‘enhancer’) with TCGA ATAC-seq open region peaks in the left panel. Right panel shown bar plot of overlaps between cancer-specific enhancer peaks with cancer-specific TCGA ATAC-seq open region peaks. **C.** Pairwise t-test p-values for H3K27Ac average intensity signal comparison between cancer types, corresponding to Figure 2D. **D.** Box plot showing mRNA expression levels for AXIN2 in cancer cell lines and matched normal samples demonstrating colon cancer-specific high expression of AXIN2. **E.** Box plot showing gene expression level of AXIN2 in all TCGA tumors and matched normals. X-axis showing different cancer types, Y-axis showing the log_2_(TPM+1) value. **F-G**. Genome browser views of ChIP-Seq tracks for H3K27Ac mark showing enhancers nearby CCND1 gene(**F**) and β-catenin gene(**G**) for each cancer type and matched normal. **H-I**. Genome browser view of tumor samples H3K27Ac ChIP-Seq signal track surrounding the CCND1 gene(**H**) and the β-catenin gene(**I**). **J-K**. Hi-ChIP plot show enhancer-promoter interaction loops for CCND1(**J**) and β-catenin(**K**) as derived from the H3K27Ac Hi-ChIP data in SW620 cells. **L-M.** Box plot showing mRNA expression levels for CCND1(**L**) and β-catenin(**M**) in cancer cell lines and matched normal samples demonstrating cancer unique high expression of CCND1 and β-catenin. In (**D-E**) and (**L-M**), the bottom and the top rectangles indicate the first quartile (Q1) and third quartile (Q3), respectively. The horizontal lines in the middle signify the median (Q2), and the vertical lines that extend from the top and the bottom of the plot indicate the maximum and minimum values, respectively.

**Extended Data Fig.3:**
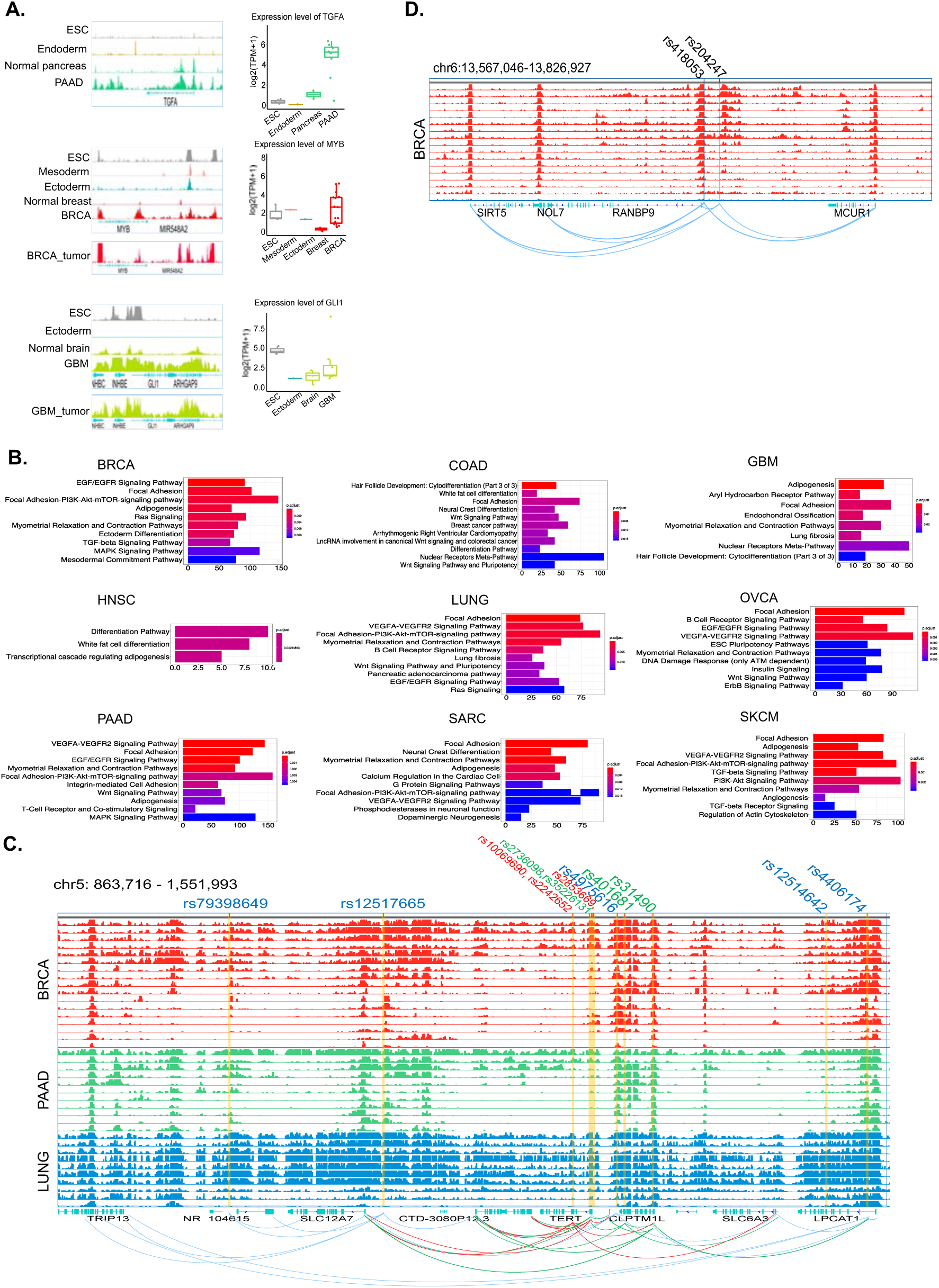
Cancer cells require cancer- and tissue-specific enhancer reprogramming during cancer development. **A**. Genome browser views of ChIP-Seq tracks for H3K27Ac mark nearby TGFα in pancreatic cancer (top), MYB in breast cancer (middle), and GLI1 in GBM (bottom) showing *de novo* and/or memory enhancer gains. Box plot on the right show gene expression levels for corresponding genes. The bottom and the top rectangles indicate the first quartile (Q1) and third quartile (Q3), respectively. The horizontal lines in the middle signify the median (Q2), and the vertical lines that extend from the top and the bottom of the plot indicate the maximum and minimum values, respectively. **B**. Bar plot showing significantly enriched pathways for genes marked by regained memory enhancers in each cancer type from ESCs during development. X-axis represents gene ratio, and colors represents adjusted p values. **C-D**. Genome browser views of ChIP-Seq tracks for H3K27Ac mark showing overlap of enhancers with important SNPs derived from various GWAS studies (shown on top of the figure). Top panel (**C**) that shows TERT gene and H3K27ac tracks from breast cancer (red), pancreatic cancer (green) and lung cancer (blue) cell lines. Whereas bottom panel (**D**) shows similar tracks nearby genes SIRT5, MCUR1 and NOL7 and overlap with breast cancer-unique SNPs. Each panel also shows interaction loops derived from HiChIP data.

**Extended Data Fig.4:**
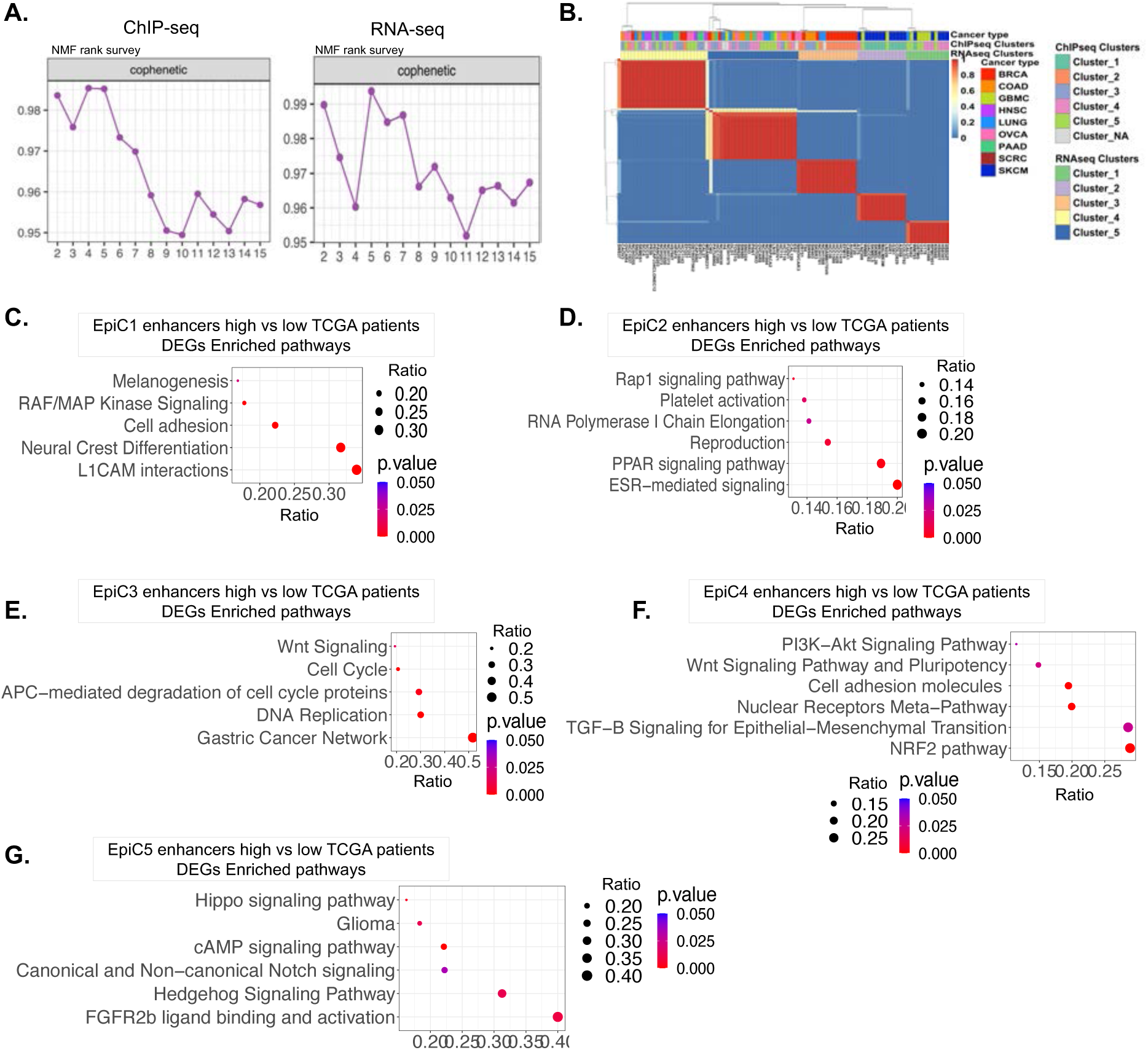
Clustering of cancer cells based on enhancer patterns. **A.** NMF clustering cophenetic scores for determining K by cluster stability in ChIP-seq (left) and RNA-seq (right) data. **B.** Heatmap of NMF clustering for RNA-seq data from cancer cell lines used in this study and overlap with cancer type and cluster annotations on top. **C-G**. Dot plot showing significantly enriched pathways for upregulated genes in top 50 patients (whose samples were profiles by the TCGA ATAC-Seq) vs bottom 50 patients based on enhancer overlap scores from EpiC1 (**C**), EpiC2 (**D**), EpiC3 (**E**), EpiC4 (**F**) and EpiC5 (**G**). Dot size represents gene ratio, and colors represent adjusted p-values.

**Extended Data Fig.5:**
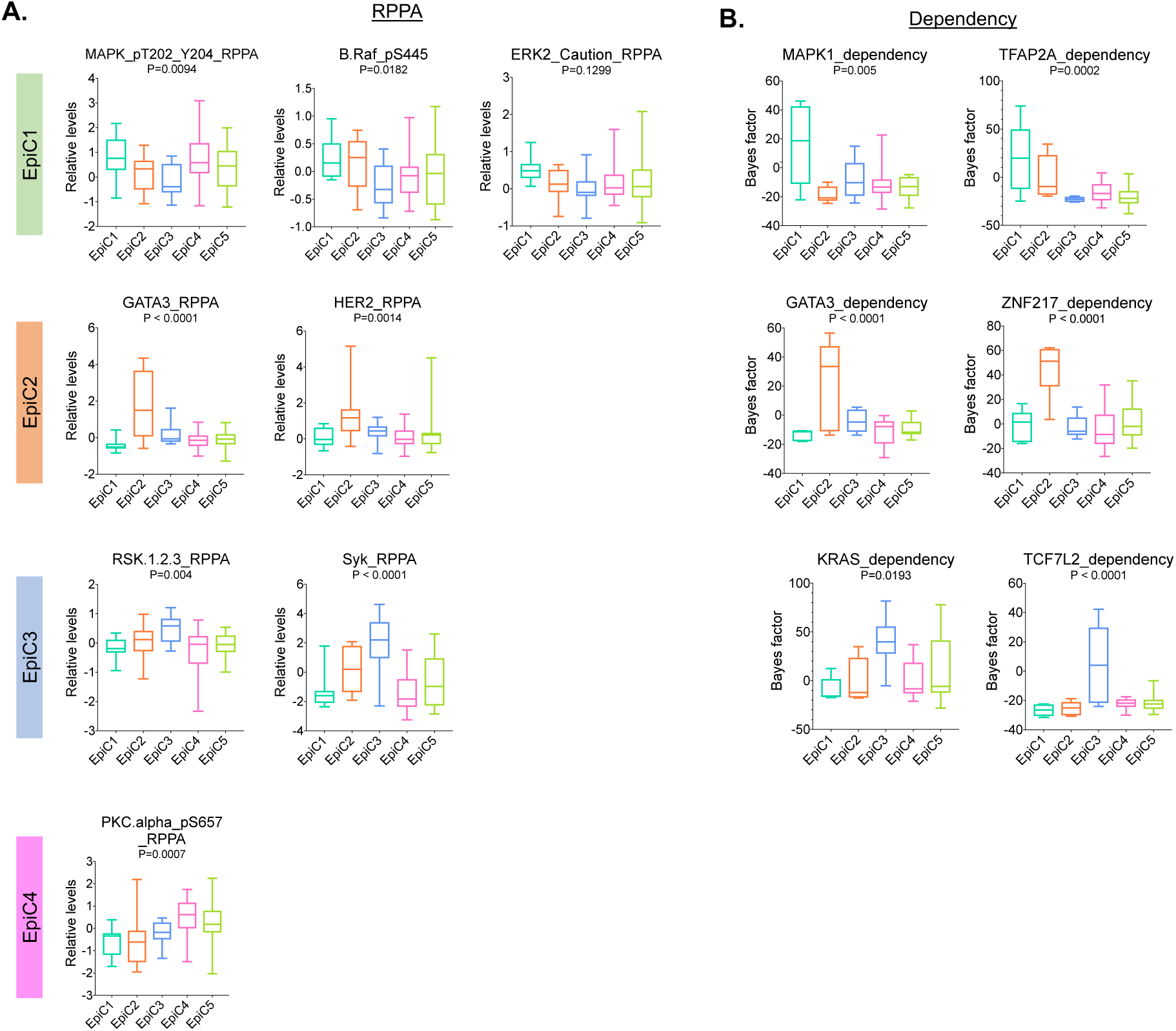
EpiC clusters specific protein activation and gene dependency. **A**. Box plot showing relative levels for EpiC-specific proteins from CCLE RPPA dataset. Colors represents the EpiC clusters shown above the plot. P-values represent one-way ANOVA comparison between 5 EpiC clusters. **B**. Box plot showing Bayes factor for EpiC-specific dependency genes from AVANA dataset. Colors represents the EpiC clusters shown above the plot. P-values represent one-way ANOVA comparison between 5 EpiC clusters. In (**A-B**), the bottom and the top rectangles indicate the first quartile (Q1) and third quartile (Q3), respectively. The horizontal lines in the middle signify the median (Q2), and the vertical lines that extend from the top and the bottom of the plot indicate the maximum and minimum values, respectively.

**Extended Data Fig.6:**
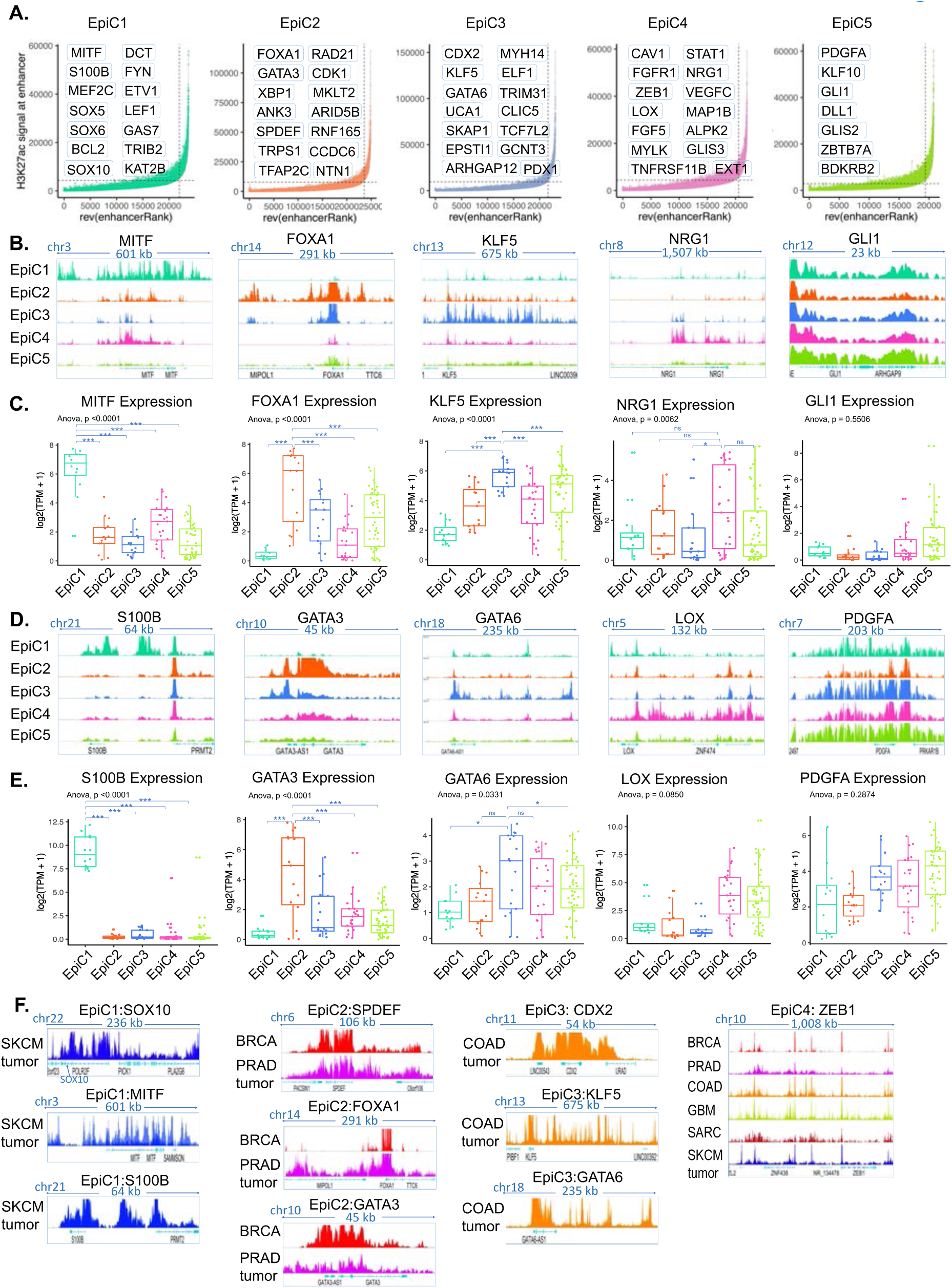
Super-enhancers regulates distinct key oncogene activities in each cluster. **A.** Inflection plot indicating super-enhancers (SEs) identified in each EpiC cluster. **B** and **D**. Genome browser view (IGV) showing composite ChIP-Seq track for H3K27Ac surrounding EpiC-specific TF genes: MITF and S100B for EpiC1, FOXA1 and GATA3 for EpiC2, KLF5 and GATA6 for EpiC3, NRG1 and LOX for EpiC4, and GLI1 and PGDFA for EpiC5. Colors represents the EpiC clusters shown above the plot. **C** and **E**. Box plot showing the gene expression level of genes targeted by EpiC-specific super-enhancers as shown in panel B and D. Each dot represents single sample. P-values represent one-way ANOVA comparison between 5 EpiC clusters. Unpaired t-test p-values were calculated for specific EpiC vs others. ns indicates non-significant. Asterisk denotes *p < 0.05, ** p < 0.001, *** p < 0.0001. The bottom and the top rectangles indicate the first quartile (Q1) and third quartile (Q3), respectively. The horizontal lines in the middle signify the median (Q2), and the vertical lines that extend from the top and the bottom of the plot indicate the maximum and minimum values, respectively. **F.** Genome browser view of composite ChIP-Seq tracks for tumor H3K27ac surrounding representative EpiC-specific TF genes: SOX10, MITF and S100B for EpiC1, SPDEF, FOXA1 and GATA3 for EpiC2, CDX2, KLF5 and GATA6 for EpiC3, and ZEB1 for EpiC4. Colors represents the cancer types shown above the plot.

**Extended Data Fig.7:**
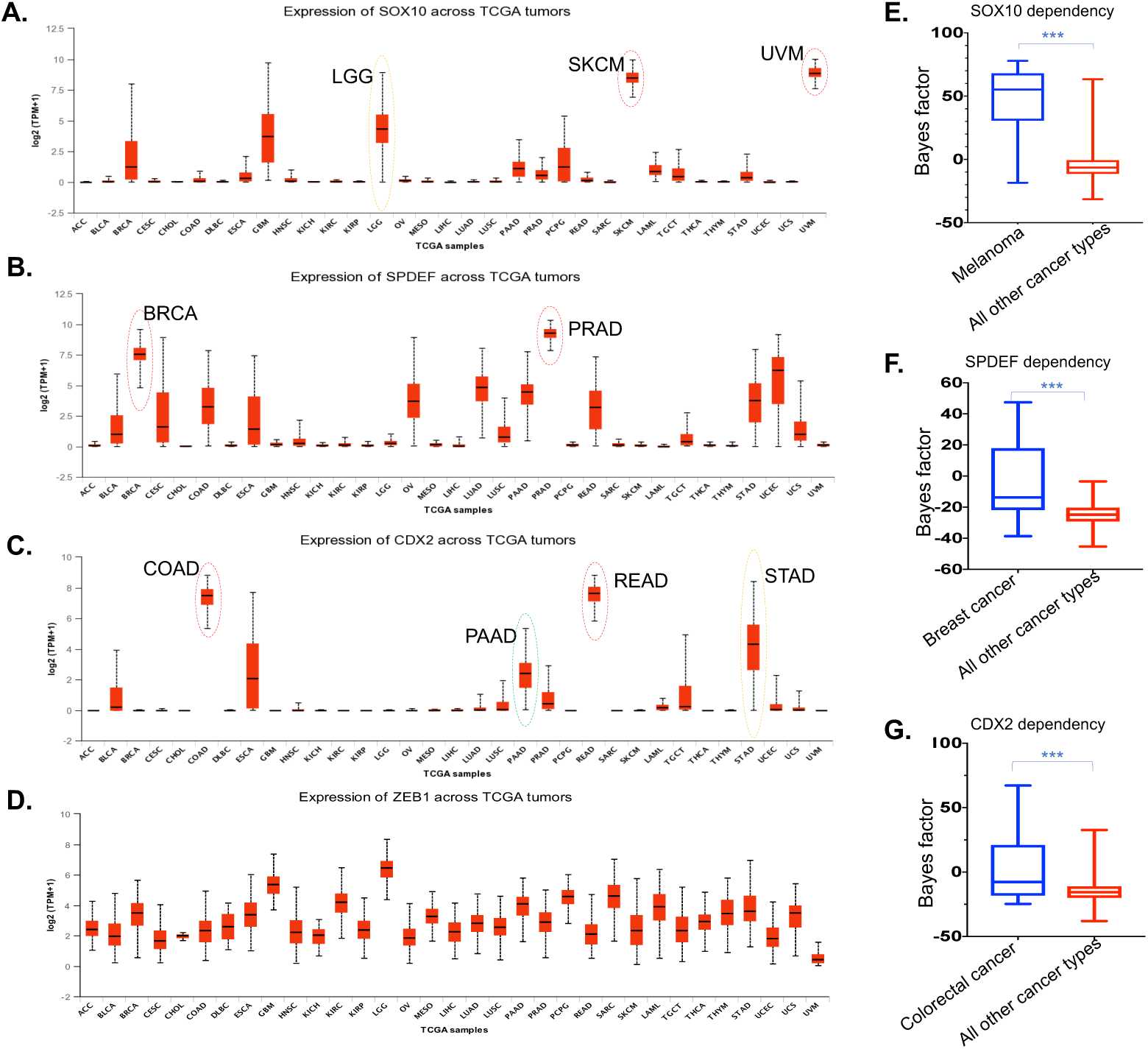
Core TFs as regulators of EpiC-specific enhancer patterns. **A-D**. Box plot showing gene expression levels for SOX10 (**A**), SPDEF (**B**), CDX2 (**C**) and ZEB1 (**D**) in the TCGA tumors. X-axis showing different cancer types, Y-axis showing the log_2_(TPM+1) value. **E-G**. Box plots comparing Bayes factor (as a proxy for dependency derived from AVANA dataset) for SOX10 between Melanoma versus all other cancer types (**E**), for SPDEF between BRCA versus all other cancer types (**F**) and for CDX2 between colorectal cancer versus all other cancer types (**G**). The p values were determined using two-tailed Student’s t test. Asterisk denotes ***p < 0.001. In (**A-G**), the bottom and the top rectangles indicate the first quartile (Q1) and third quartile (Q3), respectively. The horizontal lines in the middle signify the median (Q2), and the vertical lines that extend from the top and the bottom of the plot indicate the maximum and minimum values, respectively.

**Extended Data Fig.8:**
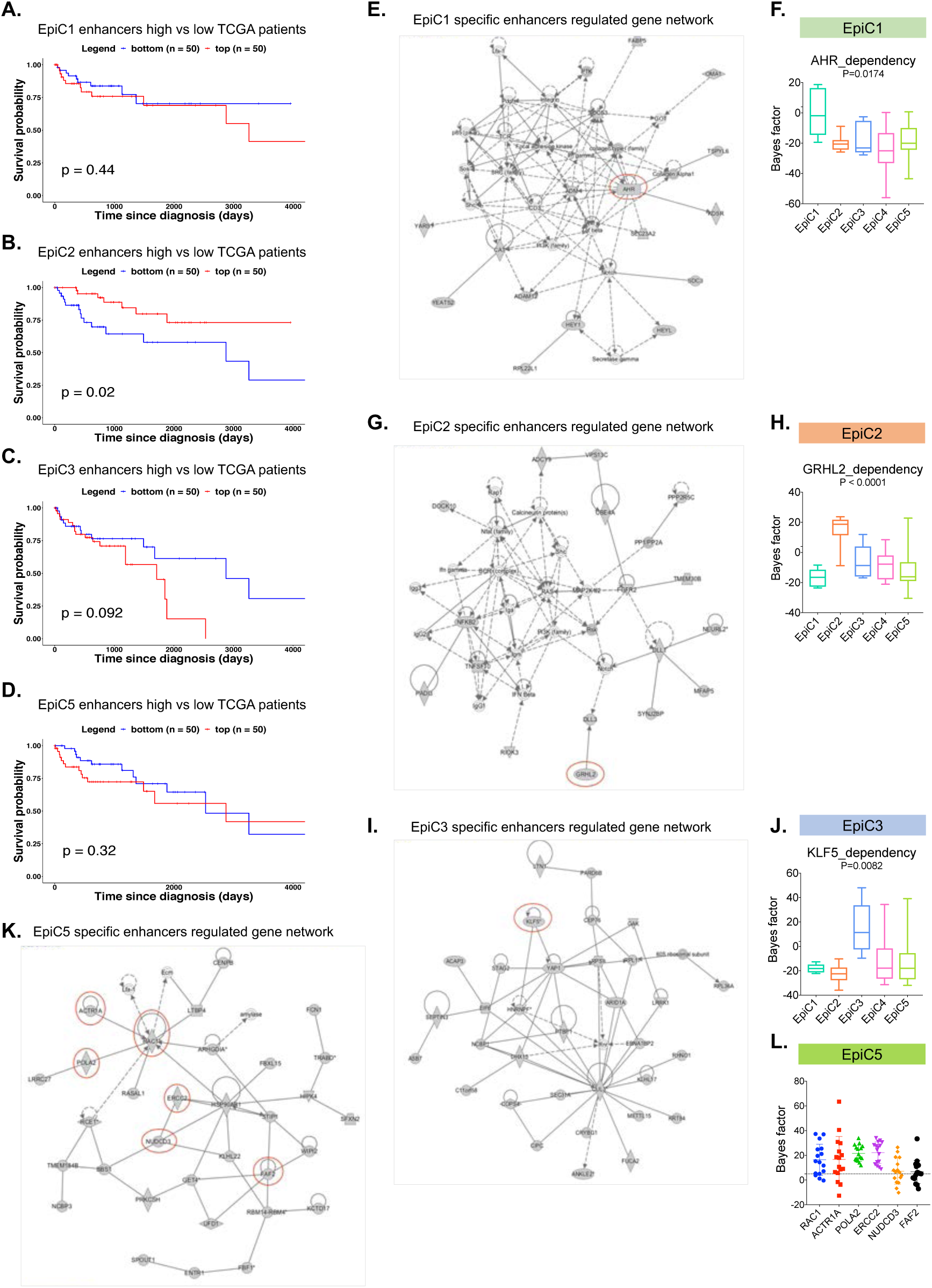
EpiC cluster specific enhancers regulatory networks. **A-D**. Kaplan-Meier analysis showing overall survival of patients (whose samples were profiles by the TCGA ATAC-Seq) with top 50 and bottom 50 normalized overlapping scores from EpiC1 (**D**), EpiC3 (**E**) and EpiC5 (**F**) overlap. p-value is derived from log rank test. **E, G, I, K**. IPA gene network analysis based on the EpiC1(**E**), EpiC2(**G**), EpiC3(**I**) and EpiC5(**K**) specific enhancer annotated genes. Red circled genes have cluster specific dependency in EpiC1(**E**), EpiC2(**G**) and EpiC3(**I**) or dependencies in EpiC5(**K**) cell lines. **F, H, J.** Box plot showing Bayes factor for EpiC1(**F**), EpiC2(**H**), EpiC3(**J**) - specific dependency genes from AVANA dataset. Colors represents the EpiC clusters shown above the plot. P-values represent one-way ANOVA comparison between 5 EpiC clusters. The bottom and the top rectangles indicate the first quartile (Q1) and third quartile (Q3), respectively. The horizontal lines in the middle signify the median (Q2), and the vertical lines that extend from the top and the bottom of the plot indicate the maximum and minimum values, respectively. **L**. Dot plot showing Bayes factor for the genes from the panel K network for EpiC5 cell lines.

